# Mass spectrometry reveals novel features of tubulin polyglutamylation in the flagellum of *Trypanosoma brucei*

**DOI:** 10.1101/2025.02.06.636787

**Authors:** Marija Nisavic, Thibault Chaze, Serge Bonnefoy, Carsten Janke, Philippe Bastin, Mariette Matondo, Julia Chamot-Rooke

## Abstract

Tubulin polyglutamylation is a prominent feature of eukaryotic cilia and flagella. In the protist parasite *Trypanosoma brucei*, nine different tubulin tyrosine ligase-like (TTLL) enzymes are potentially involved in tubulin glutamylation. Given this enzymatic diversity, it is important to qualitatively and quantitatively characterize tubulin polyglutamylation, in order to understand the trypanosome tubulin code and pave the way for functional assignments of different TTLLs. Robust mass spectrometry (MS) based proteomics approaches for analysis of posttranslational modifications have been developed in recent years, but none to study polyglutamylation. We therefore optimized a nanoLC-MS/MS pipeline, from sample preparation to data analysis, using synthetic polyglutamylated peptides for quantification. Applied to the flagellum of *T. brucei,* our approach enabled the detection and quantification of C-terminal tubulin peptides with up to eleven supplementary glutamates on α-, and five on β-tubulin. In addition to the known E445 on α- and E435 on β-, a novel glutamylation site of β-tubulin was discovered at residue E438. Furthermore, our data revealed an increase in enzymatic detyrosination with increasing length of the glutamate chains, especially for α-tubulin. This indicates crosstalk between the modifications and different rates of detyrosination of the two tubulin types. Our results complement the existing knowledge of the tubulin code in *T. brucei* and introduce an efficient analytical pipeline for the characterization of polyglutamylated proteins.

## INTRODUCTION

Cilia and flagella are widely conserved among eukaryotes where they perform motility, sensory or morphogenesis functions. They are composed of an axoneme, a cylindrical structure made of 9 doublet microtubules that surrounds a central microtubule pair in most motile cilia. The tubulins that compose axonemal microtubules are subject to extensive post-translational modifications (PTMs) which are expected to alter their biochemical properties, contributing to microtubule stability and interaction with other partners such as microtubule-associated proteins (MAPs) or molecular motors. This ensemble of modifications is now referred to as the tubulin code^1^ by analogy with the histone code. One prominent modification of flagellar tubulin is polyglutamylation. This PTM involves the addition of one or more glutamate residues to the gamma-carboxyl-group of a glutamate present in the primary sequence of C-terminal tails of both α- and β-tubulin, hence creating a branched peptide chain^2^. Glutamate residues are added by enzymes of the tubulin-tyrosine ligase-like (TTLL) family^3^ and can be removed by cytosolic carboxypeptidases^4^. Deciphering exact (poly)glutamylation patterns of tubulins and performing their quantification is challenging due to: 1) the existence of multiple genes encoding for α- and β-tubulins which differ in their C-terminal sequences, potentially leading to many tubulin proteoforms 2) the frequent presence of glycylation^5^, another competing C-terminal tubulin PTM, introduced by specific TTLLs^6^ 3) the presence of several glutamate residues in the tubulin tail offering multiple sites for glutamylation and 4) the highly acidic/hydrophilic properties of polyglutamylated peptides that complicate their analysis by classical shotgun proteomics approaches.

The flagellated protist *Trypanosoma brucei* is well known for causing sleeping sickness in Africa. In recent years, it has also emerged as a potent model organism to study cilia and flagella bringing useful insights to the comprehension of several ciliopathies^7–9^ . Trypanosomes are fascinating organisms to study since they contain a mixture of universal features but also of unique characteristics, presumably linked to their divergent position in the eukaryotic phylogeny. *T. brucei* possesses a single flagellum that is involved in cell motility but also acts as a guide for cell morphogenesis^10, 11^ and an anchor for parasite attachment to the epithelium of the salivary glands during tsetse fly infection^12^. Its length varies from 3 to 30 µm according to the life cycle stage^13, 14^. The so-called procyclic stage corresponds to the parasites developing in the midgut of the tsetse fly that can be easily cultivated and manipulated in the laboratory. Its flagellum measures about 20 µm and its composition has been established by proteomic studies^15, 16^ and a large-scale gene tagging project^17^. *T. brucei* is ideally suited to study tubulin glutamylation since it contains only one gene for α- and one for β-tubulin, albeit repeated multiple times in tandem^18^. Both α- and β-tubulin possess a final tyrosine that can be removed after microtubule assembly, increasing the diversity of tubulin proteoforms^19–21^. Importantly, glycylation seems absent in this organism, as revealed by western blot using antibodies targeting glycylated tubulin^22^, further confirmed by mass spectrometry (MS)^19^ and by the lack of tubulin glycylase genes within the TTLL family^23^. This particular feature should therefore simplify the analysis of the C-terminal modified tails by MS.

One way to study polyglutamylation is immunofluorescence labelling. Using the GT335 monoclonal antibody that detects the glutamylation branching point^24^, or polyE polyclonal antiserum that recognizes polyglutamate side chains of a minimum of three glutamate residues^25^, a strong signal on microtubules present in the subpellicular corset and the flagellar axoneme of *T. brucei*^26^ was found. However, although immunofluorescence can provide quantitative information, it cannot distinguish between residue-specific branching points or lateral chains that are more than three residues long in case of G335 or polyE, respectively. Approaches leading to more accurate results, if possible quantitative, are required. Mass spectrometry is therefore an attractive option to fulfil these goals.

The analysis of *T. brucei* tubulins by MS has been performed for the first time in 1997, using a combination of Matrix-Assisted Laser Desorption Ionization Time of Flight (MALDI-TOF) MS in negative mode and Edman sequencing^19^. For α-tubulin, peptides were obtained using an SDS-PAGE separation of cytoskeletal proteins and subsequent in-gel LysC digestion. Cyanogen bromide (CNBr) was used to generate peptides from β-tubulin. This combination of techniques led to the identification of modified peptides with 1-15 glutamate residues on glutamate 445 (E445) for α-tubulin and 1-6 glutamates on E435 for β-tubulin. Glutamylation of mammalian tubulins has also been extensively studied between 1990 and 2000 and specific MS approaches developed to this aim^27–29^. Since then, and despite considerable advances in proteomics over the last two decades both in term of instrumentation and bioinformatics, little progress has been made in the characterization of tubulin glutamylation^30^. The most recent studies, based on nanoLC-MS/MS experiments, led to the identification of only triglutamylated peptides for *Drosophila* tubulins^31^, and up to 7 glutamic acid residues for bovine samples^32, 33^, while their quantification remains a challenge due to the existence of multiple modified peptide variants.

The trypanosome genome project led to the identification of 9 candidate enzymes for tubulin polyglutamylation belonging to the TTLL family: TTLL1, TTLL4 (TTLL4A, TTLL4B & TTLL4C), TTLL6 (TTLL6A & TTLL6B), TTLL9 and TTLL12 (TTLL12A & TTL12B)^23, 26^. In agreement with the experimental absence of glycylation, homologs of the glycylases TTLL3, TTLL8 and TTLL10 are missing from *T. brucei*, as well as from the genome of all kinetoplastid species analyzed. RNAi-mediated knockdown of individual *TTLL* genes did not reveal particular phenotypes, perhaps because of redundancies between different enzymes, or low knockdown efficiency due to low abundance of target transcripts. However, combined targeting of TTLL4A and TTLL6B reduced the GT335 signal, yet without affecting the polyE signal, and led to subtle perturbations of the parasite cell cycle^26^. In a separate study, knockdown of TTLL6A or TTLL12B reduced the intensity of both GT335 and polyE signals on western blot and was accompanied by several phenotypes affecting cytoskeletal architecture and cell motility^34^, demonstrating the importance of tubulin polyglutamylation for trypanosome cell biology. Very recently, deletion of the *TTLL1* gene was reported, revealing a significant contribution to the organization of cytoplasmic microtubules at the posterior pole of the parasite^35^. So far, however, a direct impact of tubulin glutamylation on flagellum structure remains to be demonstrated. The first step toward this understanding is to achieve the most complete characterization of both α- and β-tubulin polyglutamylation, including quantification. The development of an optimized LC-MS/MS method to achieve this goal is described in this work. Our method allowed us to gain insight into the diversity of the trypanosome flagellum tubulin code, but also expands the methodological toolbox for its characterization and future functional assignments of TTLL enzymes.

## MATERIAL AND METHODS

### Trypanosome cell culture and flagellum purification

Wild-type trypanosomes (strain 427) of the procyclic stage were grown in culture in SDM- 79 medium containing 10% foetal calf serum and hemin. Cells were picked up at mid-log concentration (about 10^7^ mL^-1^) and their flagella were purified as described previously ^36^.

Briefly, pelleted cells were washed twice in phosphate-buffered. After centrifugation at 2.400 rpm, pelleted cells were gently resuspended and lysed for 2mn at room temperature in 0.1M PIPES pH6.9, 1mM MgSO4, 2mM EGTA containing 1% Nonidet NP40 substitute (IGEPAL CA-630) with protease inhibitors (Complete EDTA free cocktail (Roche) and Turbo Nuclease (Jena Biosciences) to digest DNA/RNA and prevent aggregation of flagella. After centrifugation 15,000 rpm for 30 seconds, cytoskeleton pellets were treated twice more in lysis buffer. Selective purification of flagella was subsequently achieved after depolymerization of the pellicular microtubules using high-salt concentration (1M Sodium Chloride). After centrifugation 15.000rpm for 30 seconds, flagellum pellet were treated twice more with 1M Sodium Chloride, washed quickly in de-ionized water and stored at -80°C prior mass-spectrometry experiments. After both detergent and high salt treatments, sample quality was evaluated by light microscopy.

### Sample preparation and digestion

Flagellar proteins were re-suspended in 100 µM of 8 M guanidine hydrochloride (GuHCl), containing 10 mM tris(2-carboxyethyl) phosphine (TCEP). The samples were heated at 95°C for 5 min and spun down to remove any undissolved debris. Protein concentration was checked using either Pierce 660 nm protein assay or by absorbance measurement at 280 nm using a nanodrop device. Samples were alkylated using iodoacetamide (20 mM final concentration) for 30 min in the dark, at room temperature.

### In-gel digestion

Twenty micrograms of flagellar proteins were separated using SDS-PAGE and subsequently stained with Coomassie blue. The corresponding protein band was excised and subjected to trypsin or thermolysin digestion. Each gel band was cut and washed multiple times in a solution containing 50 mM ammonium bicarbonate (ABC) and acetonitrile (1:1) for 15 minutes at 37 °C. Disulfide bonds were reduced using 10 mM DTT (43815 - Sigma, St Louis, Missouri, USA), and cysteine residues were alkylated with 55 mM chloroacetamide. Trypsin digestion was carried out overnight at 37 °C in 50 mM ABC. Thermolysin digestion was carried out overnight at 65 °C under agitation. Resulting peptides were extracted from the gel through two incubations in a solution containing 50 mM ABC, acetonitrile (ACN), and formic acid (FA) in a ratio of 50:50:0.5 for 15 minutes at 37 °C. After evaporating the ACN in a Speed-Vac, the resulting peptides were desalted using a stage-tip packed with C18 Empore discs and eluted with a solution consisting of 80% ACN and 0.1% FA. Peptides were dried and stored at -20°C until further use.

### In-solution digestion

Samples were alkylated using iodoacetamide (20 mM final concentration) for 30 min in the dark, at room temperature. To reduce the guanidine concentration. to below 1 M, 900 µL of 50 mM Tris pH 8.0 was added. Proteins were digested overnight (ON) under agitation using either trypsin (enzyme-to-substrate ratio of 1/50 ratio, at 37°C under agitation) or thermolysin (enzyme-to-substrate ratio of 1/10 ratio, supplemented with 0.5 mM Ca^2+^ final, at 65 °C). The digestion was stopped by adding trifluoroacetic acid (TFA) to a final concentration of 1%. Obtained peptides were desalted using a Sep-Pak C18 (50 mg) strategy according to manufacturer’s instructions. Briefly, C18 phase was wetted with pure methanol, activated using 80% acetonitrile (ACN), 0.1% FA and equilibrated in 0.1% FA. The loaded samples were washed with 0.1% FA, and the peptides were eluted form the phase with 50% ACN, 0.1% FA. Desalted peptides were dried and kept at -20 °C until further use.

### Synthetic peptides

For quantitation purposes, a series of peptide standards resembling the thermolysin-released C-terminal tubulin peptides (LEKDYEEVGAESADMDGEEDVEE(Y) for α- and IEEEGEFDEEEQ(Y) for β-tubulin) bearing 0, 1, 3, 5 or 10 supplementary glutamates in case of α- or 0, 1, 3 and 5 in case of β-tubulin were synthetized by PSL GmbH (Heidelberg, Germany), both in tyrosinated and detyrosinated form. Supplementary glutamates were introduced to the primary sequence in a linear fashion at position E445 of α- and position E435 of β-tubulin. Peptides were prepared as an equimolar mixture and run using the same LC-MS/MS method (see below) as the flagella samples. A total of 500 fmol of peptides, or 225 fmol spiked in 210 ng of flasdh digest (Promega) to check for matrix effect, were injected for analyses. All samples were analyzed in triplicates.

### Enrichment of polyglutamylated C-terminal peptides

#### Strong Cation-eXchange chromatography (SCX)

Peptide samples were resuspended in 0.1% acetic acid (AcOH). Fractionation was conducted using a stage tipping strategy as described ^37^. Briefly, 4 discs from Empore SCX phase were stacked in a 200 µL tip. The phase was wetted in methanol, activated in 80% ACN, 0.1% AcOH, rinsed with 500 mM ammonium acetate (AmAc), 20% ACN, 0.5% AcOH, and then rinsed with 0.5% AcOH alone. The loaded samples were washed with 0.5% AcOH, and fractionation was performed using serial elution with increasing salt buffer concentrations. The first elution was conducted using 50 mM AmAc, 20% ACN, 0.5% AcOH, the second elution using 100 mM AmAc, 20% ACN, 0.5% AcOH, and the final elution using 500 mM AmAc, 20% ACN, 0.5% AcOH. For each eluted fraction, peptides were dried and further desalted using an Agilent Bravo robot and C18 tips following the manufacturer’s instructions. Each C18 eluate was dried and stored until MS analysis.

#### Hydrophilic Interaction Liquid Chromatography (HILIC)

HILIC fractionation was conducted using a stage tipping strategy with PolyHA beads (PolyLC, 5µm, 300A) packed into a 200µl tip sealed with a C8 disc (14 gauge). Peptides were resuspended and loaded into buffer A (80% ACN, 50mM ammonium formate pH 3.0). The HILIC beads were conditioned using 5% ACN, 50mM ammonium formate, rinsed, and washed after loading with buffer A. Retained peptides were eluted in serial steps with decreasing concentrations of ACN (70% ACN, 50 mM ammonium formate; 50% ACN, 50 mM ammonium formate; 20% ACN, 50 mM ammonium formate; 10% ACN, 50 mM ammonium formate). The fractionated peptides were dried and stored until further analysis by LC-MS/MS.

#### Titanium dioxide (TiO_2_)

Titanium dioxide (TiO2) fractionation was performed using a stage tipping strategy with TiO2 beads (Sachtopore NP beads, 5 µM, 300 Å – Huntsman) packed into a 200 µL tip sealed with a C8 disc (14 gauge). A slurry was prepared with 30% ACN and 0.1% TFA. Peptides were resuspended in buffer A (80% ACN, 0.1% TFA) and loaded onto the beads, followed by rinsing with buffer. A second wash was conducted using 50% ACN and 0.1% TFA. Elution was carried out in two steps: first, using 10% NH4OH to elute acidic peptides (in a tube buffered with 20% FA), and second, using 80% ACN and 2% FA to elute peptides retained in the C8 disc. Enriched peptides from both elutions were combined and lyophilized. The peptides were stored at -80°C until further analysis by LC-MS/MS.

#### Strong anion exchange chromatography (SAX)

SAX fractionation was performed using 9 discs of SAX material (Empore Anion-SR) stacked in 200 µL tips. SAX buffers containing 20 mM acetic, 20 mM phosphoric, and 20 mM boric acid were prepared at different pH values (pH 11.3, 9.5, 7.1, 3.5 and 2) by the addition of NaOH. NaCl was added to the final elution buffer (pH 2) to a concentration of 250 mM. SAX phase was wetted in methanol, activated in 1M NaOH and equilibrated with SAX buffer of pH 11.3. The samples were re-suspended in, loaded and washed with SAX buffer of pH 11.3. The elution was performed using the prepared buffers of descending pH. All fractions were dried and further desalted using C18 stage tips in a standard fashion. Each C18 eluate was speed vac dried and kept at -20°C until MS analysis.

### LC-MS/MS

#### Non-enriched & SCX, HILIC and TiO_2_ enriched peptides

Peptides were reconstituted in a solution containing 2% acetonitrile (ACN) and 0.1% formic acid (FA). LC-MS/MS analysis of enriched polyglutamylated peptides was conducted using an Orbitrap Q Exactive Plus mass spectrometer (Thermo Fisher Scientific, Bremen) coupled to an EASY-nLC 1200 system (Thermo Fisher Scientific). A custom-made column packed with C18 material (30 cm capillary column picotip silica emitter tip, 75 μm diameter filled with 1.9 μm Reprosil-Pur Basic C18- HD resin, Dr. Maisch GmbH, Ammerbuch-Entringen, Germany) was employed for peptide separation. Prior to sample loading, the column was equilibrated, and peptides were loaded at a pressure of 900 bars in buffer A (0.1% FA). Peptide separation was achieved at a flow rate of 250 nL/min. A gradient of solvent B (ACN with 0.1% FA) was used for peptide elution: starting from 3% B, increasing to 7% over 8 minutes, then to 23% over 95 minutes, and finally to 45% over 45 minutes. The total chromatographic run time, including a high ACN level step and column regeneration, was 132 minutes. The column temperature was maintained at 60°C throughout the analysis. Mass spectra were acquired in data-dependent acquisition mode using XCalibur software (v2.2, Thermo Fisher Scientific, Bremen). MS scans were performed at a resolution of 70,000, while MS/MS scans (fixed first mass 100 m/z) were acquired at a resolution of 17,500. The AGC (Automatic Gain Control) target and maximum injection time for both survey and MS/MS scans were set to 3E6 and 20 ms, and 1E6 and 60 ms, respectively. An automatic selection of the 10 most intense precursor ions (Top 10) with a dynamic exclusion of 45 seconds was employed. The isolation window was set to 1.6 m/z, and a fixed normalized collision energy of 28 was used for higher-energy collisional dissociation (HCD) fragmentation. An underfill ratio of 1.0% corresponding to an intensity threshold of 1.7E5 was applied. Unassigned precursor ion charge states, as well as charge states of 1, 7, 8, and >8, were rejected, and peptide matching was disabled.

#### SAX enriched peptides

Peptides were reconstituted in 2% ACN, 0.1% FA. The LC-MS/MS analysis was performed on an Orbitrap Eclipse mass spectrometer (Thermo Fisher Scientific, Bremen) coupled to Vanquish Neo Liquid Chromatography (Thermo Fisher Scientific). An Easy-Spray PepMap Neo column was used for peptide separation (C18, 75 cm, 75 μm x 750 mm (Thermo Fisher Scientific). The column was equilibrated, and the samples were loaded in solvent A (0.1% FA). The peptides were separated and eluted at 300 nL/min using an increasing gradient of solvent B (80% ACN, 0.1% FA) from 2% to 31% in 70 min, 31% to 60% in 25 min (total length of the chromatographic run was 113 min including high ACN level steps and column regeneration). Mass spectra were acquired in data-dependent acquisition mode using the XCalibur 2.2 software (Thermo Fisher Scientific, Bremen) with automatic switching between MS and MS/MS scans using a 3 s cycle time method. MS spectra were recorded at a resolution of 60,000 (at m/z 400) with a target value of 4 × 10^5^ ions over a maximum injection time of 50 ms. The scan range was set from 300 to 1700 m/z. Peptide fragmentation was performed using CID (collision energy 35%, activation time 10 ms and activation Q 0.25). All MS/MS were recorded in the Orbitrap cell at a resolution of 15,000 (at m/z 400). Intensity threshold for ions selection was set at 1 × 10^5^ ions with charge exclusion of z=1 and z>7. Isolation window was set at 1.6 Th. Dynamic exclusion was employed within 25s. As the first flagella biological replicate was used for optimization, it was recorded as a single measurement. The second biological replicate was recorded as a technical triplicate.

### Data analysis

Raw data were searched against *Trypanosoma brucei brucei* database (downloaded on 25.04.2023) using MetaMorpheus software (v. 1.0.2 and 1.0.5)^38, 39^. The spectra were first calibrated at both MS1 and MS2 level using default parameters. GPTMD was used to search for multiple glutamylations on E residues. The following G-PTM-D settings were used: protease = thermolysin; maximum missed cleavages = 5; minimum peptide length = 7; maximum peptide length = 25; dissociation type = CID (For Orbitrap Tribrid Eclipse data) and HCD (for Orbitrap Q Exactive Plus data), initiator methionine behaviour = Variable; max modification isoforms = 1024; fixed modifications = Carbamidomethyl on C, Carbamidomethyl on U; variable modifications = Oxidation on M. G-PTM-D modifications concerned all ‘common biological’ and custom glutamylation with up to fifteen supplementary glutamates on E. Precursor and fragment tolerance were set to 5 ppm and 20 ppm, respectively. FlashLFQ was employed at 5 ppm peakfind tolerance^40^. All other parameters for the Search task were used as default with match between run (MBR) option on. All identifications with score ≥12 and FDR of 1% on both peptide and protein level were considered for Eclipse data and score ≥6 for Q-Exactive data to take into account the difference in sensitivity of both instruments.

For quantification purposes, two prevalent series of C-terminal peptides bearing all detected glutamylated variants were selected (LEKDYEEVGAESADMDGEEDVEE(Y) for α- and IEEEGEFDEEEQ(Y) for β-tubulin). Due to ambiguity in site-specific information of some polyglutamylated peptide variants, label-free quantitation was performed per number of supplementary glutamates (Tables S1-S8). The quantitation is based on intensities from “all quantified peaks” file generated by Metamorpheus search. This is done to ensure that all glutamylated peptides are included in the quantitation, irrespective if the “exact” modification site was determined by the software or not. Furthermore, since supplementary glutamates were introduced as variable modifications and fragmentation of the lateral chain is not considered, this approach minimizes potential errors in software site-specific assignments that could arise as a result of overlapping fragment masses.

Presence/absence of terminal tyrosine and number of supplementary glutamates introduce changes in the sequences of the C-terminal tubulin peptides that can reflect on peptide ionization efficiency which, in turn, can result in erroneous quantitation. Data on equimolar mixture of synthetic peptides were recorded in order to check for these differences. It was searched against tubulin database using the same Metamorpheus settings used to search the flagella samples. As the MS intensity response curves for 500 fmol and 225 fmol (in yeast digest) injections showed a very similar profile (Figure S1), 500 fmol injection was chosen for further calculations. Intensity values for peptides with 2, 4, 6, 7, 8 and 9 glutamates for α- or 2 and 4 glutamates for β-tubulin were estimated via a linear regression between neighbouring points for each, tyrosinated and detyrosinated variants of both proteins (Figures S2 and S3). A series of correction factors were introduced based on ratio of intensity between detyrosinated mono- glutamylated α-tubulin sequence, selected as a reference as it was the most prevalent species in the flagella samples, and intensity of each peptide in the mixture (Table S9). Although variations in calculated ratios can be expected in between different experiments, this correction serves as an approximation, to estimate the extent of the influence of differences in sequence-based ionization efficiency on peptide label-free quantities (LFQ).

## Data Availability

Proteomics data have been deposited in the PRIDE^41^ database via the ProteomeXchange Consortium (http://www.ebi.ac.uk/pride) with the dataset identifier PXD059627.

## RESULTS AND DISCUSSION

### Purification of trypanosome flagella

Flagella were purified from exponentially growing wild-type trypanosomes of the procyclic stage using an optimized protocol based on detergent-extraction of the whole cytoskeleton (Figure 1A), followed by incubation with 1M NaCl to depolymerise the subpellicular microtubules^36^, with the addition of DNase to prevent aggregate formation. Their quality was checked by light microscopy which confirmed the absence of cell debris or other remnants of the cytoskeleton (Figure 1B). One-dimension SDS gel electrophoresis showed the expected abundance of the tubulin bands (TUB), as well as the major paraflagellar rod (PFR) proteins (Figure 1C).

**Figure 1.**
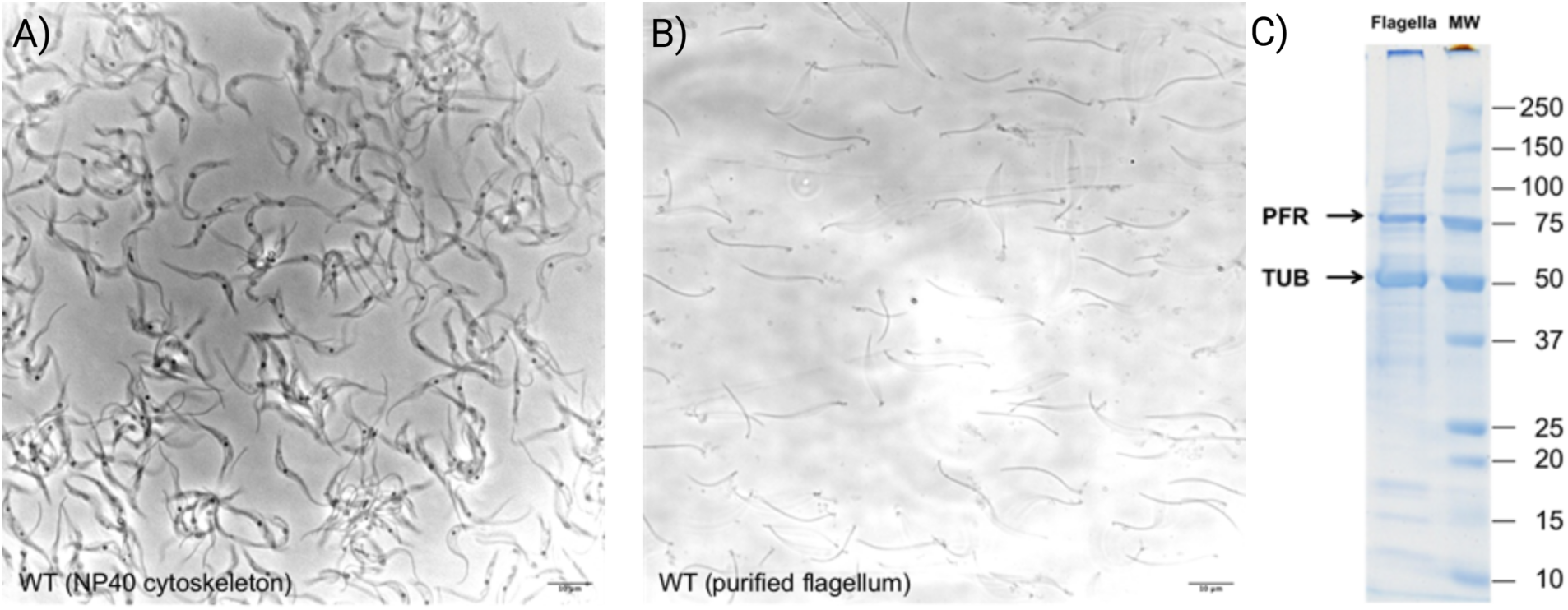
Quality control of purified flagella. Light microscopy analysis of detergent-extracted cytoskeletons (A) and of purified flagella (B), confirming the absence of debris or other contaminants. (C) SDS-PAGE analysis of the flagellar fraction confirms the expected abundance of tubulin (TUB) and paraflagellar rod (PFR) proteins.

### Different strategies for enrichment of polyglutamylated peptides

We first analyzed the tryptic digest of purified flagella using nanoLC-MS/MS on a Q-Exactive mass spectrometer (non-enriched samples). The results confirmed that trypsin was not the most suitable enzyme for identifying tubulin C-termini, with only α-tubulin leading to peptides in this region (data not shown). For β-tubulin, the expected C-terminal tryptic peptides are probably too large to be detected. We therefore searched for the most appropriate enzyme and identified thermolysin as a good candidate, potentially leading to C-terminal peptides for both α- and β-tubulins in a mass range compatible with the nanoLC-MS/MS analysis. The predicted cleavage sites of α- and β-tubulin C-terminus for various enzymes including trypsin, AspN and thermolysin are reported in Figure S1.

The analysis of flagella samples digested with thermolysin confirmed that it was more suitable than trypsin, but also revealed that the detected glutamylated peptides were of low abundance. To overcome this limitation, we decided to test several enrichment strategies to achieve the most comprehensive characterization of polyglutamylated peptides. We evaluated both in-gel and in-solution digestion with thermolysin, coupled with enrichment using chromatographic techniques including HILIC^42^, TiO2^43^, SCX, and SAX^44^. These methods have previously demonstrated efficacy in analyzing post-translationally modified peptides, particularly those with acidic properties. For negatively charged peptides, SCX is considered as a “negative” enrichment method, which means that the expected peptides are in the flow through of the first fraction.

The results depicted in the supplemental Table S1a and Table S1b shows the value of using thermolysin digestion to identify C-terminal glutamylated peptides for both α- and β-tubulin. For α-tubulin, in-gel thermolysin digestion combined with HILIC and SCX enrichment produced various glutamylated peptide sequences, with 1-3 extra glutamates mainly located on E445 (Table S1a), confirming the results previously obtained^19^ . The precise localization of these extra glutamates from MS/MS data is complicated for two main reasons. The first one is the presence of consecutive glutamate residues in the backbone sequence (such as E445 and E446 for α-tubulin or E438, E439 and E440 for β-tubulin) that leads to an ambiguity if the sequence coverage is not 100% in this region. The second one is the possibility of combining backbone cleavage with side chain ones (in particular for long stretches of glutamates), which cannot be differentiated. For these reasons, Metamorpheus sometimes proposes two different assignments with the same score. For samples digested in liquid, only the HILIC enrichment led to useful information with also 1-3 extra glutamates, again found on E445.

For β-tubulin, a smaller number of modified peptides were obtained, with a combination of 1-4 glutamates distributed on E435 but also surprisingly on E438, a modification site which has never been described previously (Table S1b). For instance, HILIC enrichment leads exclusively to di- and tri-glutamylated peptides at E438. In general, the number of extra glutamates was higher for β-tubulin compared to α-tubulin, with less singly modified peptides and more doubly and triply modified ones.

To improve the depth of analysis, we decided both to test a SAX fractionation and analyze our enriched samples with an Eclipse Tribrid Orbitrap (Figure 2). Since each glutamate adds a negative charge to C-terminal peptides, SAX fractionation can represent an attractive option.

**Figure 2.**
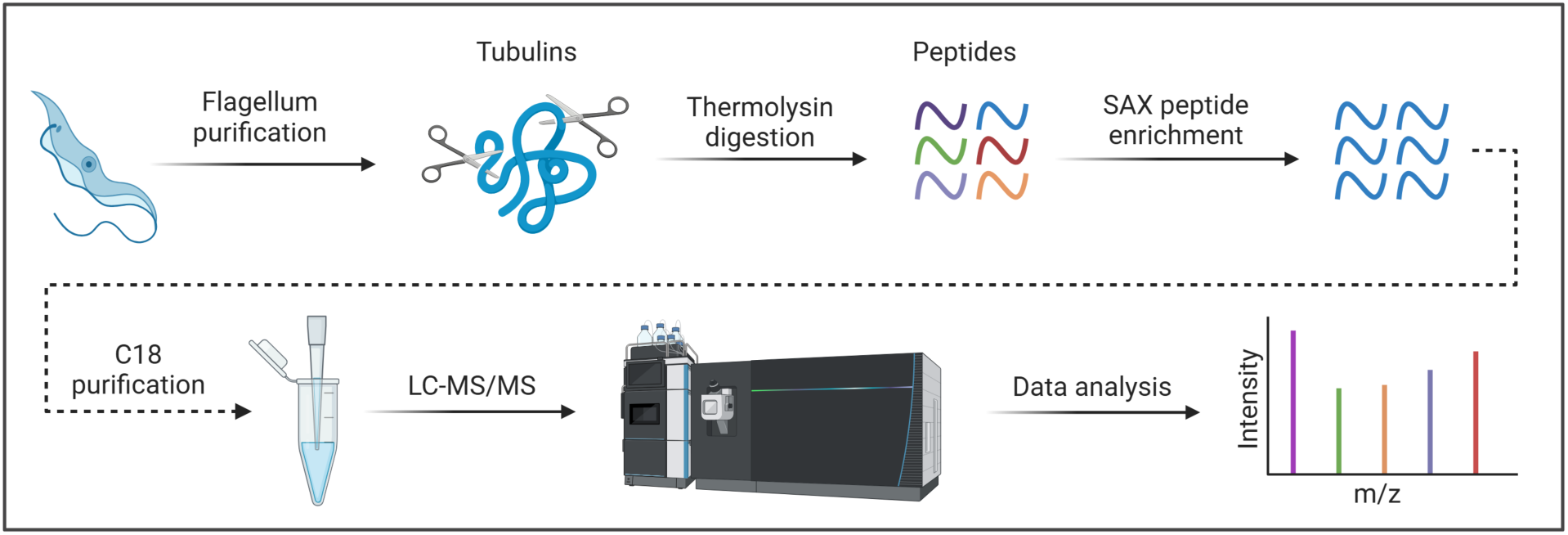
Preferred workflow for characterization of tubulin glutamylation used in this study.

A first biological replicate of *T. brucei* flagella was used for the optimization of several experimental parameters (LC gradient, fragmentation conditions,…). The improved sensitivity of the whole pipeline enabled the detection of a significantly higher number of glutamylated peptides than previously observed with a maximum of 11 additional glutamates for α- tubulin (Table S2a) and 5 for β-tubulin (Table S2b). As observed for the other enrichment methods, E445 was identified as the most prevalent modification site for α-tubulin (examples of annotated MS/MS spectra in Figure S8-S11)^19^. Three additional candidate modification sites (E446 as previously observed, but also E449 or E450) were also identified by the software (Figure S12-S13). However, since each glutamylation was introduced as a variable modification, fragmentation of the side chain that happens in parallel with the fragmentation of the peptide backbone is not taken into consideration in the assignment. Side chain fragmentation is clearly visible in MS/MS spectra of e.g., long-chain modifications where a series of unassigned high m/z fragment ions with mass difference corresponding to a single glutamate can be observed, often accompanied by a loss of water (Figure S17). This simultaneous fragmentation can result in overlapping fragment ions that can be difficult or impossible to distinguish which, in some cases, introduces ambiguity in obtaining site-specific information. Since no evidence for the additional candidate sites were previously found using Edman sequencing^19^, E445 is most likely the only modification site on α-tubulin decorated with polyglutamate chains of varying lengths (from 1 to 11E). As previously observed with other enrichment approaches, all longer polyglutamylation chains were found exclusively in a detyrosinated form (Figure 3, left) (examples of annotated MS/MS spectra in Figure S14-S17).

**Figure 3.**
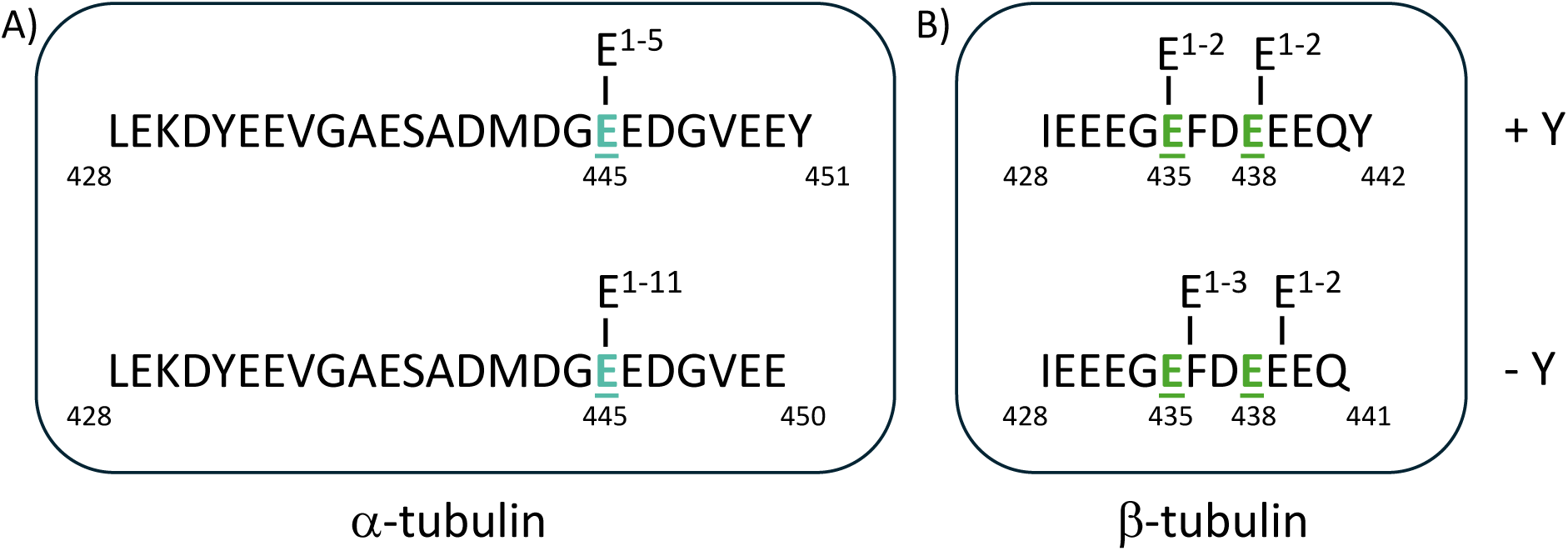
Tyrosinated (+Y) and detyrosinated (-Y) A) α-tubulin and B) β-tubulin C-terminal peptide polyglutamylation patterns in the flagellum of *T. brucei*.

For β-tubulin, the close analysis of MS/MS spectra enabled the unambiguous determination of two distinct modification sites: E435 which was the first previously discovered and E438 as newly identified in our HILIC or TiO2 data (Table S1b). In contrast to α-tubulin, long side chain modifications (E>5) were not found. The longest detected polyglutamylation (5E), was found to modify exclusively detyrosinated β-tubulin occupying both E435 and E438 sites, carrying 3 and 2 glutamates, respectively (Figure 3, right). All β-tubulin and more than 95% of α-tubulin peptides were identified solely in their glutamylated forms, confirming extensive tubulin glutamylation in the trypanosome flagellum.

In order to confirm the obtained results, we analysed a second biological replicate of purified flagella (in technical triplicate). Although most identified peptides overlap with the single measurement of the first biological replicate, we failed to detect α-tubulin peptides bearing more than seven additional glutamates in this experiment. Apart from possible biological variations, one explanation could be the poor ionizability of acidic peptides in positive ion mode. Due to this, the risk of poor reproducibility between experiments and even technical replicates is therefore quite high, as even minor variations in sample processing, chromatographic separation or spray stability can influence the analysis. Further, the stochastic nature of data dependent acquisition (DDA) methods that rely on the fragmentation of a preselected number of the most abundant precursor ions (“TopN”) results in only partially reproducible LC-MS/MS runs^45, 46^, although the match-between-run option that we enabled here partially solves this issue. As previously observed, all glutamylated peptides were commonly detected with one additional charge to what is theoretically expected^32, 33^. Examples of annotated MS/MS spectra are shown in Figure S18-S20 for tyrosinated β-tubulin peptides and Figure S21-S25 for detyrosinated ones.

### Quantification of polyglutamylated peptide variants and their detyrosination levels

Quantifying individual polyglutamylated peptide variants can be a considerable challenge, bearing in mind that multiple factors apart from abundance govern the MS response. These include the nature of the analyte, matrix, chromatographic and ionization conditions^47, 48^. Even though in case of *T. brucei*, the absence of different tubulin isotypes and of polyglycylation simplifies the analysis, it is important to note that each modification to the sequence can cause significant changes in the peptide ionization efficiency. Here, these changes stem from a variable number of supplementary glutamates and the presence or absence of the final tyrosine. To gain insight into the extent of these changes and their influence on LFQ, we used synthetic peptides bearing a varying number of supplementary glutamates each in a tyrosinated and detyrosinated form. We analyzed an equimolar mixture of these synthetic peptides directly or spiked into a tryptic yeast digest to mimic a complex mixture, in the same experimental conditions as our *T. brucei* flagella sample. As expected (Figure S1-S3), a difference in the ionization efficiency of these peptides could be observed since the equimolar amount injected does not lead to peaks of the same intensities for the ionized species. Correction factors were thus calculated from the observed differences in intensities (Table S5). The objective of this correction factor is to allow for the global quantification of polyglutamylated peptides based on the MS1 peak intensities. For peptides lacking a synthetic counterpart, an average of the correction factors obtained for the two neighbouring peptides was used. All correction factors were then applied to the flagella datasets. Calculated relative abundances of each detected peptide variant belonging to detyrosinated and tyrosinated α- tubulin for the first biological replicate are shown in Figure 4 before (A-B) and after (C-D) correction.

**Figure 4.**
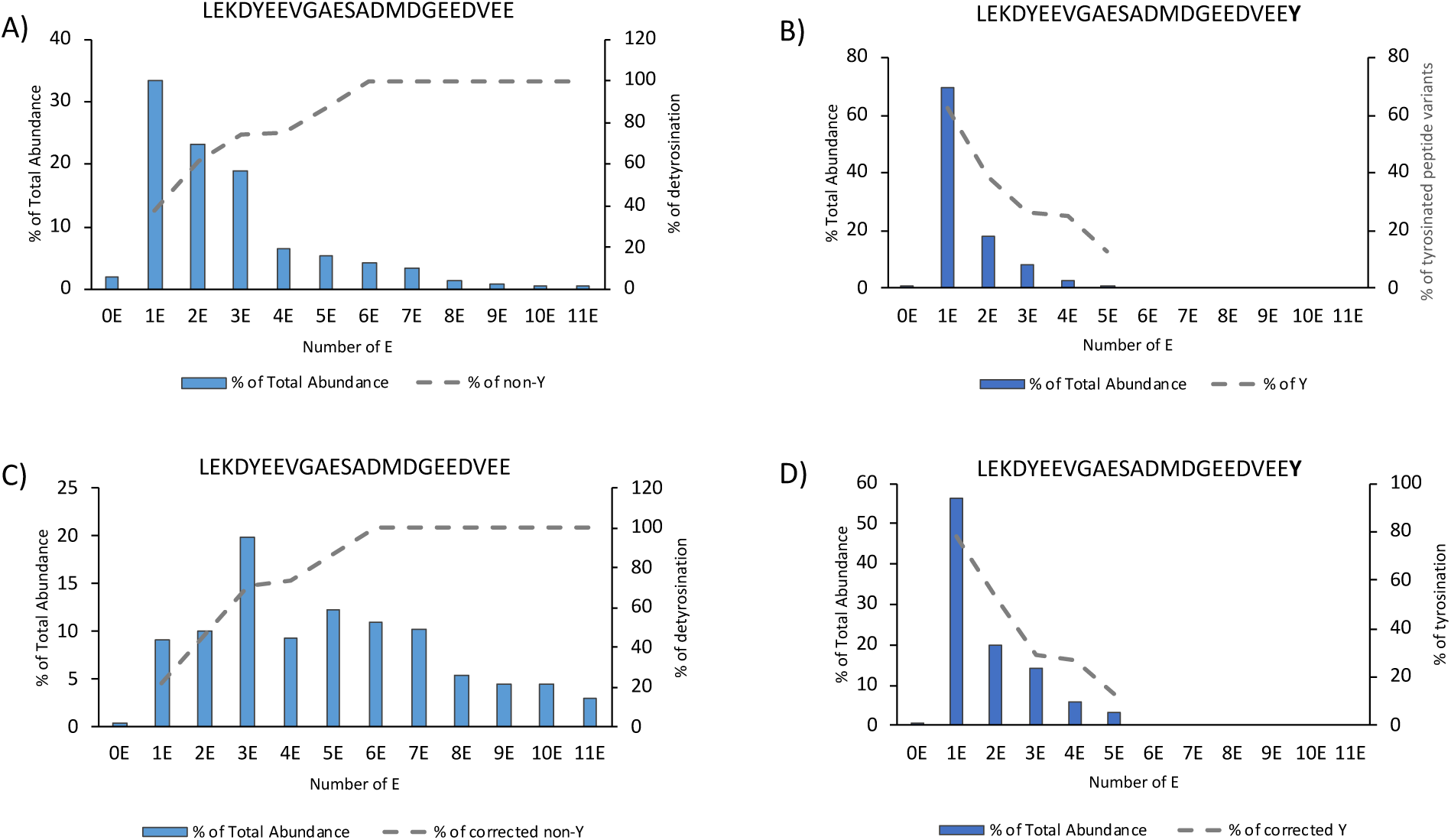
Quantification of alpha-tubulin C-terminal peptide variants identified in the first biological replicate: A) detyrosinated before correction B) tyrosinated before correction C) detyrosinated after correction D) tyrosinated after correction. Dashed lines represent the variation in detyrosination (A and C) or tyrosination (B and D) with increasing the number of supplementary glutamates.

It can be noted that the relative abundances change upon correction. In particular, the intensities of long-chain species, that are underrepresented in the non-corrected data, increase. This is expected, as the longest chains with multiple glutamate residues (up to 10) introduce the largest differences to the sequence, exhibit the lowest ionization efficiency and are, therefore, the least reflective of LFQ abundance. The analysis of α-tubulin for the second biological replicate is shown in Figure 5 (top). Here, as polyglutamylations with more than seven supplementary glutamates were not detected, applied correction has lesser influence on the overall distribution amongst peptide variants comparing to the first replicate. Nevertheless, the general trend is similar: the detyrosinated species are more heavily glutamylated and more evenly distributed comparing to tyrosinated forms where the mono-glutamylated variant is by far the most abundant (Figure 5, A-B).

**Figure 5.**
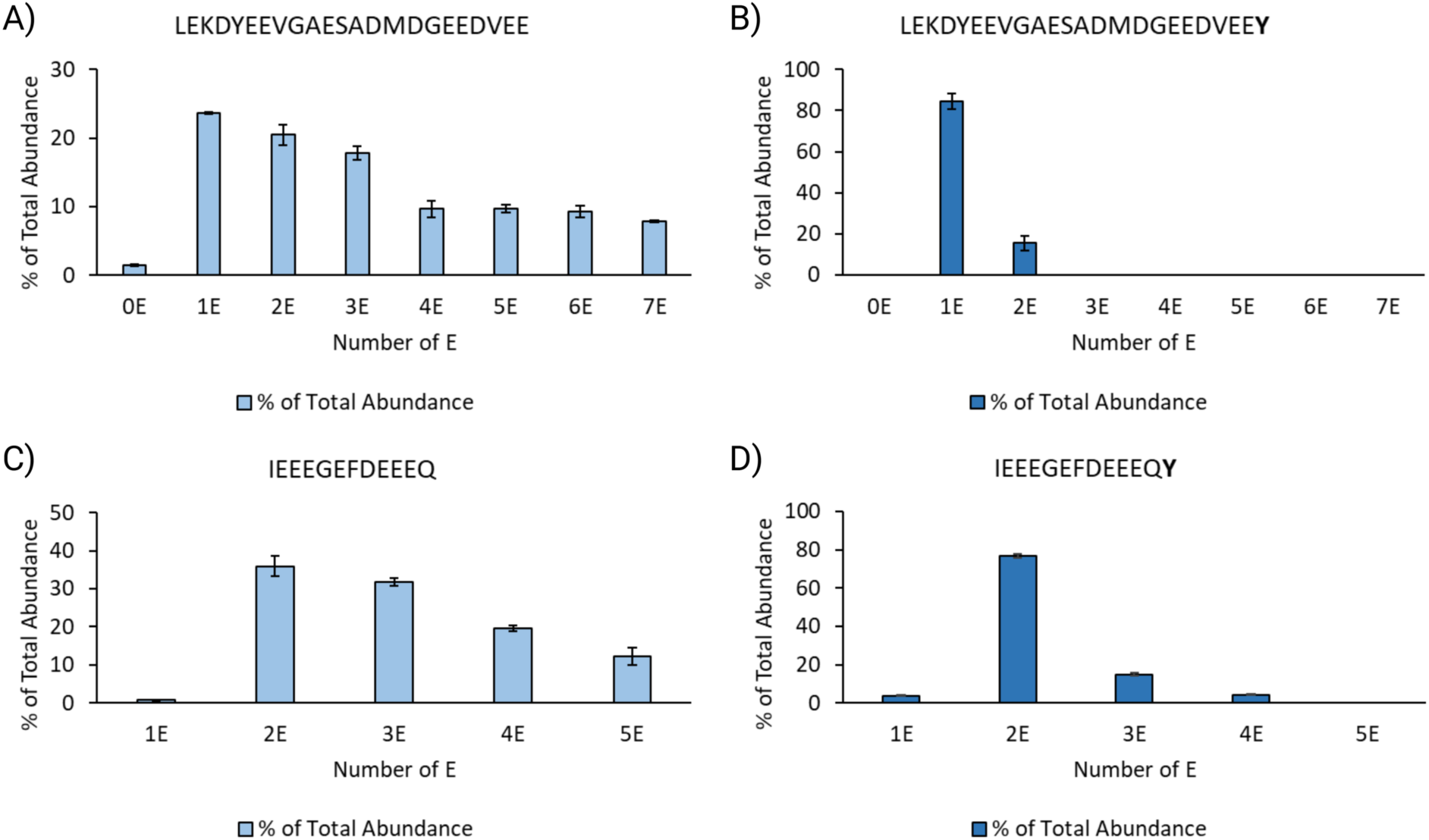
Quantification of C-terminal peptide variants of detyrosinated (A) and tyrosinated (B) α-tubulin and detyrosinated (C) and tyrosinated (D) β-tubulin (from the second biological replicate).

Analysis of β-tubulin showed very similar profiles between the first (Figure S5) and the second biological replicate (Figure 5 C-D), with correction having negligible influence on the distribution of abundances. Here, di- and tri-glutamylated variants are the most abundant in the detyrosinated form of the protein, while the di-glutamylated variant dominates the tyrosinated protein. In both cases, as well as in the case of α-tubulin (particularly non-corrected data), the obtained quantitative estimates correlate well with the results previously obtained by MALDI-TOF in negative mode^19^.

As previously highlighted, most long-chain polyglutamylation carrying peptides are detected in detyrosinated form of both proteins, particularly so in case of α-tubulin. Relative abundances of each detected variant, with or without the C-terminal tyrosine, with respect to their total abundance are shown in Figure S7 for the first, or Figure 6 for the second biological replicate. These data demonstrate not only distinct and unique polyglutamylation, but also specific detyrosination profiles of α- and β-tubulin in *T. brucei,* which had not been characterized earlier.

**Figure 6.**
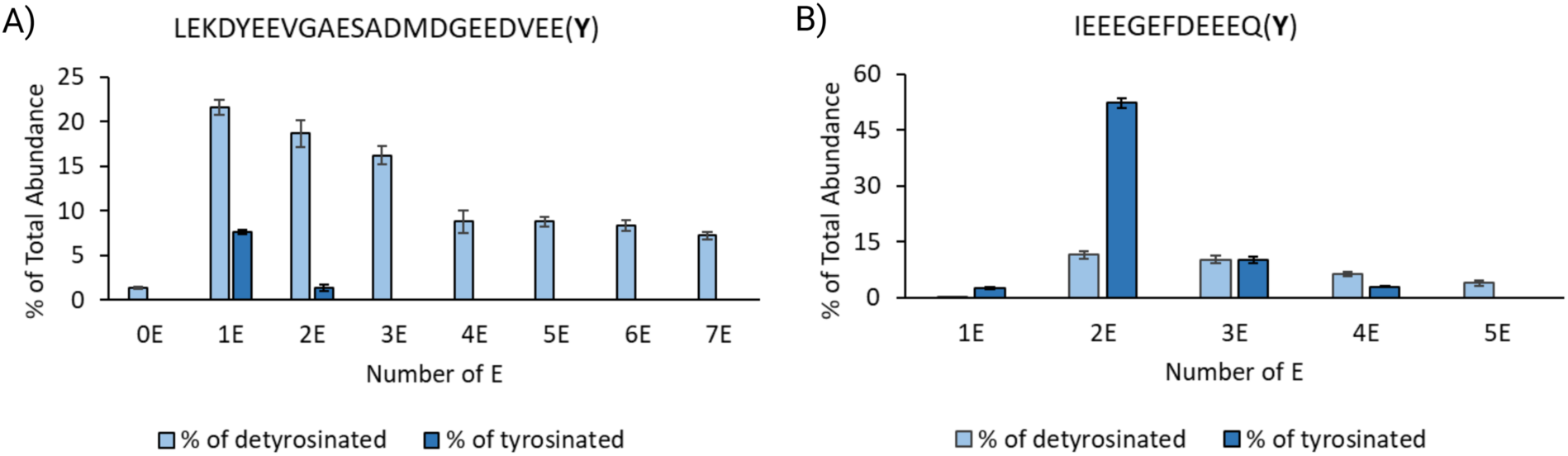
Relative abundances of detyrosinated and tyrosinated forms of C-terminal peptide variants of α- (A) and β-tubulin (B). For α-tubulin, detyrosinated forms are highly predominant although this is the opposite for β-tubulin.

Together with the estimates of the extent of total detyrosination, it can be seen that the majority of α-tubulin is found in its detyrosinated form (Fig. 6A & Fig. 7), while this does not appear to be the case for β-tubulin (Fig. 6B & Fig. 7). This clearly indicates that, despite being performed by the same enzyme^21^ the rate of detyrosination might not be equal for both proteins and appears to happen at a slower rate in case of β-tubulin.

**Figure 7.**
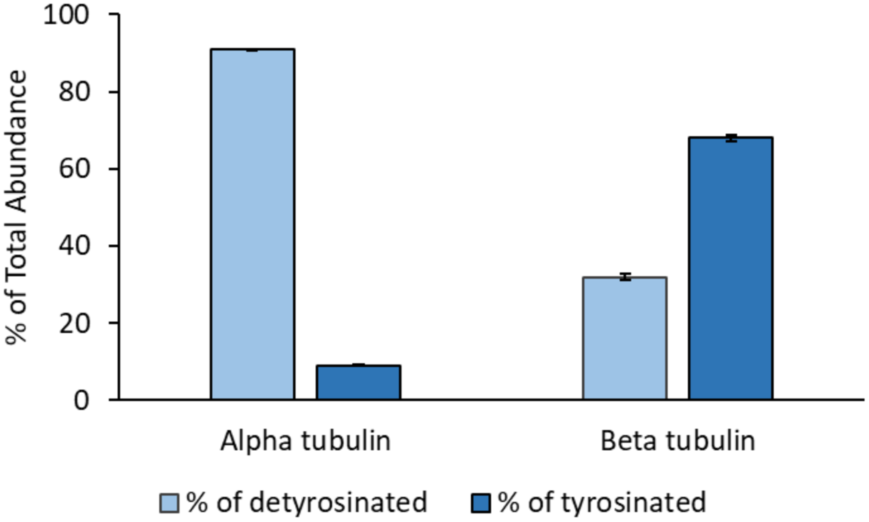
Relative abundances of detyrosinated and tyrosinated α- and β-tubulin in *T. brucei*.

### Interplay between polyglutamylation and detyrosination

In a recent study, it has been shown that polyglutamylation of α-tubulin promotes its detyrosination by increasing enzymatic activity of the tubulin tyrosine carboxypeptidase vasohibin/small vasohibin-binding protein (VASH/SVBP) complex^49^. The extent of activity enhancement was directly related to the length of the lateral glutamate chains, with a 16-fold increase observed for microtubules decorated with 10E-long lateral chains compared to the non-glutamylated ones. Lack or under-representation of the final tyrosine in long-chain polyglutamylated tubulin (Figures 5 and S8) observed in our data suggests a similar effect of tubulin polyglutamylation on *Trypanosoma* VASH homolog carboxypeptidase activity. This enzyme performs detyrosination of α- but also of β-tubulin which, in Trypanosomatid parasites, uniquely possesses a C-terminal tyrosine^21^. The differences in detyrosination and polyglutamylation profiles of α- and β-tubulin C-terminal peptide variants observed herein demonstrate that these PTMs are distinctly regulated in the two tubulin types, presumably through different activities of VASH and TTLL enzymes. Bearing in mind that polyglutamylation of β-tubulin proved to be less extensive compared to that of α-, with respect to the length of the lateral chains, reduced VASH activity and consequent reduction in detyrosination is expected and in line with our results. Indeed, human VASH2, which was demonstrated to be similar to trypanosome VASH with respect to autonomous catalytic activity^21^, possesses a positively charged surface surrounding the active site, where it binds the negatively charged tubulin tails^49^. Therefore, increased polyglutamylation could modulate this interaction, resulting in increase of VASH association or enzymatic activity. Similarly, a separate *in vitro* study showed that the human TTLL6 enzyme is 8-fold more efficient on the detyrosinated version of α-tubulin^50^. Although analogous effect could also take place in *T. brucei*, van der Laan et al. reported that VASH knockout parasites do not display detectable changes in tubulin glutamylation when using the polyE antibody in western blotting^21^. This could indicate that glutamylation might be the primary regulator of detyrosination in *T. brucei*, but not vice versa. However, mass spectrometry analysis of the tubulin tails in the VASH knockout cell line has not been reported, so discrete perturbations cannot be ruled out. Although further studies are needed to accurately elucidate this interplay in different species, crosstalk between detyrosination and glutamylation is apparent from this as well as the supporting studies, and it could be of great importance for the regulation and maintenance of the tubulin code.

## CONCLUSIONS

Using an optimized nanoLC-MS/MS strategy based on a SAX enrichment, we achieved the, so far, most comprehensive investigation of tubulin glutamylation in *T. brucei* flagella. We found the glutamylation profile of β-tubulin to be quite distinct to that of α-tubulin. As expected, E435 was frequently glutamylated but surprisingly, we discovered that E438 was almost as prominent. This position had not been reached by sequencing in the Schneider et al.^19^ study and it is, to our knowledge, the first demonstration of polyglutamylation of trypanosome β-tubulin at that position. Most detected peptides were found to carry glutamate side chains at both residues, as reflected by the high relative abundance of di-glutamylated species. Those with a single glutamate were present in weak amounts and modified only on E435, never on E438, suggesting that, in a sequence of events, E435 is the first residue to be glutamylated. Therefore, one could consider that E435 needs to be glutamylated before the enzyme modifying E438 can act.

Using polyglutamylated peptides as external standards, we showed that peptide ionization efficiencies clearly depend on the number of supplementary glutamates and presence/absence of the terminal tyrosine, allowing for correction of the relative abundance of the different modifications and revealing the abundance of long-chain polyglutamylations of α-tubulin. One could envision that herein suggested polyglutamylation profiling could be used to determine which TTLL enzymes are responsible for different modifications, something that so far relied on antibody-based experiments^26, 34, 35^. Our data also shed light on detyrosination of both α- and β-tubulin, with a more exhaustive detyrosination for α-tubulin. A clear increase in detyrosination with increasing length of polyglutamate lateral chains is observed. The detyrosination reaction takes place after the assembly of microtubules and its timing could be monitored closely using the YL1/2 antibody that recognises only the tyrosinated form of α- tubulin^51^. Knowing that the growth rate of the flagellum is around 4 µm per hour^52^, it means that detyrosination is complete in a bit more than one hour after tubulin incorporation in axonemal microtubules. Furthermore, labelling with radioactive tyrosine proved that retyrosination does not take place on flagellar microtubules, in contrast to those present in the cortex^53^. Because of sequence differences, the YL1/2 antibody cannot detect tyrosination on trypanosome β-tubulin, hence precluding similar analysis. Additional evidence for a crosstalk between detyrosination and glutamylation comes from the recently reported deletion of TTLL1. Loss of this polyglutamylase not only led to a disorganization of cytoplasmic microtubules, but also an increased level of tubulin tyrosination at the posterior tip of the cell revealed upon staining with YL1/2 antibody^35^.

Our study has revealed complex features of tubulin polyglutamylation in the trypanosome flagellum, an essential step in elucidating the tubulin code. It opens the door to global functional investigations, knowing that up to 9 candidate enzymes for initiation or elongation of glutamate side chains have been identified in the genome^26^. Initial investigations by RNAi knockdown or gene deletion of some of these enzymes have started to lift the veil on important roles of tubulin polyglutamylation in cytoskeleton assembly or cell motility^26, 34^. Further understanding will require coupling functional studies to MS analyses as reported here to evaluate the tubulin PTM landscape in flagella of mutant cells since glutamylases have been shown to compete with, or to substitute to, each other^6^.

## Supporting information

Calculations for all figures

## ASSOCIATED CONTENT

### Supporting Information

Table S1a. Glutamylated peptides identified for α-tubulin using different analytical strategies.

Table S1b. Glutamylated peptides identified for β–tubulin using different analytical strategies.

Table S2a. Glutamylated peptides identified for α-tubulin using SAX (first biological replicate).

Table S2b. Glutamylated peptides identified for β-tubulin using SAX (first biological replicate).

Table S3a. Glutamylated peptides identified for detyrosinated α-tubulin (second biological replicate).

Table S3b. Glutamylated peptides identified for tyrosinated α-tubulin (second biological replicate).

Table S4a. Glutamylated peptides identified for detyrosinated β-tubulin (second biological replicate).

Table S4b. Glutamylated peptides identified for tyrosinated β-tubulin (second biological replicate).

Table S5. List of synthetic peptides and correction factors applied to corresponding peptides (500 fmol injection).

Figure S1. Predicted cleavage sites of α- and β-tubulin C-terminus for trypsin (Tryps), AspN and thermolysin (Therm).

Figure S2. Synthetic peptide MS intensity response corresponding to alpha- (A-B) and beta- (C-D) tubulin C-terminals recorded at 500 fmol injection (left) or 225 fmol injection spiked in yeast digest (right).

Figure S3. Synthetic peptide (500 fmol injection) MS intensity response corresponding to alpha- (A-B) or beta- (C-D) tubulin C-terminals in their detyrosinated (left) and tyrosinated (right) forms.

Figure S4. Synthetic peptide (500 fmol injection) MS intensity response corresponding to A) alpha- or B) beta-tubulin C-terminal ones.

Figure S5. Quantification of β-tubulin C-terminal peptide variants identified in the first biological replicate: A) detyrosinated before correction B) tyrosinated before correction C) detyrosinated after correction D) tyrosinated after correction.

Figure S6. Relative abundances of detyrosinated and tyrosinated (poly)glutamylated peptide variants of α-tubulin A) before and C) after correction and β-tubulin B) before and D) after correction with respect to their total abundance.

Figure S7. Relative abundances of detyrosinated and tyrosinated α- and β-tubulin in the first biological replicate A) before and B) after correction.

Figure S8-S17. MS/MS spectra of wild type flagella α-tubulin peptides. Figure S18-S25. MS/MS spectra of wild type flagella β-tubulin peptides.

The Calculations-for-Figures-SI file (XLSX) gathers all calculations leading to Figures 4, 5A- 5B, 5C-5D, 6A, 6B, 7 and Figures S5A, S5B, S6A-S6B, S6C, S6D and S7.

## AUTHOR INFORMATION

The authors declare no competing financial interests.

## ACKNOWLEDGMENT

We thank Dr. Karen Druart for her help in the Metamorpheus analysis of data from SCX, HILIC and TiO2 enrichment.

