## Supplementary information for "Mass spectrometry reveals novel features of tubulin polyglutamylation in the flagellum of *Trypanosoma brucei*"

**Table S1a.** Glutamylated peptides identified for α-tubulin using different analytical strategies [Ox]: oxidized

| Sample type | Enzyme | Enrichment Method | Peptide sequence | | Number of E |
|---|---|---|---|---|---|
| | | | **Detyrosinated** | **Tyrosinated** | |
| Gel based | Trypsin | SCX | DYEEVGAESADM[ox]DGE[**2E**]EDVEE<br>OR<br>DYEEVGAESADM[ox]DGEE[**2E**]DVEE | | 2E |
| | | HILIC | NA | | |
| | | $TiO_2$ | DYEEVGAESADM[ox]DGE[**1E**]EDVEE or<br>DYEEVGAESADM[ox]DGEE[**1E**]DVEE | DYEEVGAESADM[ox]DGE[**1E**]EDVEEY or<br>DYEEVGAESADM[ox]DGEE[**1E**]DVEEY or<br>DYEEVGAESADM[ox]DGEEDVE[**1E**]EY | 1E |
| | | | DYEEVGAESADM[ox]DGE[**2E**]EDVEE or<br>DYEEVGAESADM[ox]DGEE[**2E**]DVEE | | 2E |
| | | | DYEEVGAESADM[ox]DGE[**3E**]EDVEE | | 3E |
| | Thermolysin | SCX | ADM[ox]DGE[**1E**]ED<br>ADM[ox]DGE[**1E**]EDVEE | ADMDGE[**1E**]EDVEEY<br>ADM[ox]DGE[**1E**]EDVEEY | 1E |
| | | | ADM[ox]DGE[**2E**]EDVEE | | 2E |
| | | HILIC | ADM[ox]DGE[**1E**]EDVEE | ADM[ox]DGE[**1E**]EDVEEY<br>M[ox]DGE[**1E**]EDVEEY | 1E |
| | | | ADM[ox]DGEE[**3E**]DVEE | ADM[ox]DGE[**3E**]EDVEEY<br>OR<br>ADM[ox]DGEE[**3E**]DVEEY | 3E |
| | | $TiO_2$ | NA | | |
| Liquid based | Trypsin | SCX | NA | | |
| | | HILIC | NA | | |
| | | $TiO_2$ | NA | | |
| | Thermolysin | SCX | NA | | |
| | | HILIC | ADM[ox]DGE[**1E**]ED<br>ADM[ox]DGEE[**1E**]D<br>M[ox]DGE[**1E**]EDVEE | ADM[ox]DGE[**1E**]EDVEEY<br>M[ox]DGE[**1E**]EDVEEY | 1E |
| | | | ADM[ox]DGE[**2E**]EDVEE<br>M[ox]DGE[**2E**]EDVEE | | 2E |
| | | | VGAESADM[ox]DGEE[**3E**]DVEE | | 3E |
| | | $TiO_2$ | VGAESADM[ox]DGE[1E]EDVEE | | 1E |

**Table S1b.** Glutamylated peptides identified for β–tubulin using different analytical strategies.

| Sample type | Enzyme | Enrichment Method | Peptide sequence | | Number of E |
|---|---|---|---|---|---|
| | | | **Detyrosinated** | **Tyrosinated** | |
| Gel based | Trypsin | SCX | NA | | |
| | | HILIC | NA | | |
| | | $TiO_2$ | NA | | |
| | Thermolysin | SCX | | IEEEGE[**1E**]FDEEEQY | 1E |
| | | | IEEEGEFDEE[**2E**]EQ | | 2E |
| | | | IEEEGEFDE[**3E**]EEQ | | 3E |
| | | | IEEEGEFDEE[**3E**]EQ | | |
| | | | IEEEGEFDE[**2E**]E[**2E**]EQ | | 4E |
| | | HILIC | IEEEGEFDEE[**2E**]EQ | | 2E |
| | | | IEEEGEFDEE[**3E**]EQ | | ~~3E~~ |
| | | $TiO_2$ | IEEEGEFDEE[**2E**]EQ | | 2E |
| | | | IEEEGEFDE[**3E**]EEQ | | 3E |
| Liquid based | Trypsin | SCX | NA | | |
| | | HILIC | NA | | |
| | | $TiO_2$ | NA | | |
| | Thermolysin | SCX | IEEEGE[**2E**]FDE[**1E**]EEQ | | 3E |
| | | | IEEEGEFDE[**1E**]E[**3E**]EQ | | 4E |
| | | HILIC | IEEEGE[**1E**]FDE[**1E**]EEQ | IEEEGE[**1E**]FDE[**1E**]EEQY | 2E |
| | | | IEEEGE[**2E**]FDE[**1E**]EEQ | | 3E |
| | | $TiO_2$ | IEEEGE[**1E**]FDE[**1E**]EEQ | | 2E |
| | | | IEEEGE[**2E**]FDE[**1E**]EEQ | | 3E |

**Table S2a.** Glutamylated peptides identified for α-tubulin using SAX (first biological replicate)

| Number of E | Peptide | |
|---|---|---|
| | **Detyrosinated** | **Tyrosinated** |
| 0E | LEKDYEEVGAESADMDGEEDVEE | |
| 1E | LEKDYEEVGAESADMDGE[**1E**]EDVEE | LEKDYEEVGAESADMDGE[**1E**]EDVEEY |
| 2E | LEKDYEEVGAESADMDGE[**2E**]EDVEE<br>**OR**<br>LEKDYEEVGAESADMDGE[**1E**]EDVEE[**1E**] | LEKDYEEVGAESADMDGE[**2E**]EDVEEY |
| 3E | LEKDYEEVGAESADMDGE[**3E**]EDVEE<br>**OR**<br>LEKDYEEVGAESADMDGE[**2E**]EDVEE[**1E**] | LEKDYEEVGAESADMDGE[**3E**]EDVEEY<br>**OR**<br>LEKDYEEVGAESADMDGE[**2E**]E[**1E**]DVEEY<br>**OR**<br>LEKDYEEVGAESADMDGE[**2E**]EDVEE[**1E**]Y |
| 4E | LEKDYEEVGAESADMDGE[**3E**]E[**1E**]DVEE<br>**OR**<br>LEKDYEEVGAESADMDGE[**3E**]EDVEE[**1E**] | LEKDYEEVGAESADMDGE[**2E**]E[**2E**]DVEEY<br>**OR**<br>LEKDYEEVGAESADMDGE[**3E**]EDVEE[**1E**]Y |
| 5E | LEKDYEEVGAESADMDGE[**3E**]EDVEE[**2E**]<br>**OR**<br>LEKDYEEVGAESADMDGE[**5E**]EDVEE | LEKDYEEVGAESADMDGE[**5E**]EDVEEY |
| 6E | LEKDYEEVGAESADMDGE[**6E**]EDVEE<br>**OR**<br>LEKDYEEVGAESADMDGE[**5E**]EDVEE[**1E**]<br>**OR**<br>LEKDYEEVGAESADMDGE[**5E**]EDVE[**1E**]E | |
| 7E | LEKDYEEVGAESADMDGE[**6E**]EDVEE[**1E**]<br>**OR**<br>LEKDYEEVGAESADMDGE[**7E**]EDVEE<br>**OR**<br>LEKDYEEVGAESADMDGE[**6E**]E[**1E**]DVEE | |
| 8E | LEKDYEEVGAESADMDGE[**7E**]EDVEE[**1E**]<br>**OR**<br>LEKDYEEVGAESADMDGE[**6E**]EDVEE[**2E**]<br>**OR**<br>LEKDYEEVGAESADMDGE[**7E**]EDVE[**1E**]E | |
| 9E | LEKDYEEVGAESADMDGE[**8E**]EDVEE[**1E**]<br>**OR**<br>LEKDYEEVGAESADMDGE[**7E**]EDVE[**2E**]E<br>**OR**<br>LEKDYEEVGAESADMDGE[**7E**]EDVEE[**2E**] | |
| 10E | LEKDYEEVGAESADMDGE[**9E**]EDVEE[**1E**]<br>**OR**<br>LEKDYEEVGAESADMDGE[**8E**]EDVEE[**2E**] | |
| 11E | LEKDYEEVGAESADMDGE[**8E**]EDVE[**3E**]E<br>**OR**<br>LEKDYEEVGAESADMDGE[**9E**]EDVEE[**2E**] | |

**Table S2b.** Glutamylated peptides identified for β-tubulin using SAX (first biological replicate)

| Number of E | Peptide | |
|---|---|---|
| | **Detyrosinated** | **Tyrosinated** |
| 1E | IEEEGE[**1E**]FDEEEQ | IEEEGE[**1E**]FDEEEQY |
| 2E | IEEEGE[**1E**]FDE[**1E**]EEQ | IEEEGE[**1E**]FDE[**1E**]EEQY |
| 3E | IEEEGE[**2E**]FDE[**1E**]EEQ<br>**OR**<br>IEEEGE[**1E**]FDE[**2E**]EEQ<br>**OR**<br>IEEEGE[**1E**]FDEE[**2E**]EQ | IEEEGE[**1E**]FDE[**2E**]EEQY<br>**OR**<br>IEEEGE[**2E**]FDE[**1E**]EEQY |
| 4E | IEEEGE[**2E**]FDE[**2E**]EEQ | IEEEGE[**2E**]FDE[**2E**]EEQY |
| 5E | IEEEGE[3**E**]FDE[**2E**]EEQ | |

**Table S3a.** Glutamylated peptides identified for detyrosinated α-tubulin (second biological replicate)

| Number of E | Replicate 1 | Replicate 2 | Replicate 3 |
|---|---|---|---|
| 0E | LEKDYEEVGAESADMDGEEDVEE | LEKDYEEVGAESADMDGEEDVEE | LEKDYEEVGAESADMDGEEDVEE |
| 1E | LEKDYEEVGAESADMDGE[**1E**]EDVEE | LEKDYEEVGAESADMDGE[**1E**]EDVEE | LEKDYEEVGAESADMDGE[**1E**]EDVEE |
| 2E | LEKDYEEVGAESADMDGE[**2E**]EDVEE<br>**OR**<br>LEKDYEEVGAESADMDGE[**1E**]E[**1E**]DVEE | LEKDYEEVGAESADMDGE[**2E**]EDVEE | LEKDYEEVGAESADMDGE[**2E**]EDVEE<br>**OR**<br>LEKDYEEVGAESADMDGE[**1E**]EDVEE[**1E**] |
| 3E | LEKDYEEVGAESADMDGE[**3E**]EDVEE<br>**OR**<br>LEKDYEEVGAESADMDGE[**2E**]EDVEE[**1E**] | LEKDYEEVGAESADMDGE[**3E**]EDVEE | LEKDYEEVGAESADMDGE[**3E**]EDVEE<br>**OR**<br>LEKDYEEVGAESADMDGE[**2E**]EDVEE[**1E**] |
| 4E | LEKDYEEVGAESADMDGE[**3E**]EDVEE[**1E**] | LEKDYEEVGAESADMDGE[**3E**]EDVEE[**1E**]<br>**OR**<br>LEKDYEEVGAESADMDGE[**3E**]E[**1E**]DVEE | LEKDYEEVGAESADMDGE[**3E**]EDVEE[**1E**] |
| 5E | LEKDYEEVGAESADMDGE[**5E**]EDVEE | LEKDYEEVGAESADMDGE[**5E**]EDVEE | LEKDYEEVGAESADMDGE[**5E**]EDVEE |
| 6E | LEKDYEEVGAESADMDGE[**6E**]EDVEE<br>**OR**<br>LEKDYEEVGAESADMDGE[**5E**]E[**1E**]DVEE | LEKDYEEVGAESADMDGE[**6E**]EDVEE<br>**OR**<br>LEKDYEEVGAESADMDGE[**5E**]EDVEE[**1E**] | LEKDYEEVGAESADMDGE[**5E**]EDVEE[**1E**]<br>**OR**<br>LEKDYEEVGAESADMDGE[**5E**]EDVE[**1E**]E |
| 7E | LEKDYEEVGAESADMDGE[**6E**]EDVEE[**1E**] | LEKDYEEVGAESADMDGE[**6E**]EDVEE[**1E**]<br>**OR**<br>LEKDYEEVGAESADMDGE[**6E**]EDVE[**1E**]E | LEKDYEEVGAESADMDGE[**6E**]EDVEE[**1E**] |

**Table S3b.** Glutamylated peptides identified for tyrosinated α-tubulin (second biological replicate)

| Number of E | Replicate 1 | Replicate 2 | Replicate 3 |
|---|---|---|---|
| 1E | LEKDYEEVGAESADMDGE[**1E**]EDVEEY | LEKDYEEVGAESADMDGE[**1E**]EDVEEY | LEKDYEEVGAESADMDGE[**1E**]EDVEEY |
| 2E | LEKDYEEVGAESADMDGE[**2E**]EDVEEY (MBR) | LEKDYEEVGAESADMDGE[**2E**]EDVEEY | LEKDYEEVGAESADMDGE[**2E**]EDVEEY (MBR) |
| 3E | LEKDYEEVGAESADMDGE[**3E**]EDVEEY | LEKDYEEVGAESADMDGE[**3E**]EDVEEY | LEKDYEEVGAESADMDGE[**3E**]EDVEEY |

**Table S4a.** Glutamylated peptides identified for detyrosinated β-tubulin (second biological replicate)

| **Number of E** | **Replicate 1** | **Replicate 2** | **Replicate 3** |
|---|---|---|---|
| 1E | IEEEGE[**1E**]FDEEEQ | NA | IEEEGE[**1E**]FDEEEQ |
| 2E | IEEEGE[**1E**]FDE[**1E**]EEQ | IEEEGE[**1E**]FDE[**1E**]EEQ | IEEEGE[**1E**]FDE[**1E**]EEQ |
| 3E | IEEEGE[**1E**]FDE[**2E**]EEQ<br>**OR**<br>IEEEGE[2**E**]FDE[**1E**]EEQ | IEEEGE[**2E**]FDE[**1E**]EEQ<br>**OR**<br>IEEEGE[**1E**]FDE[**2E**]EEQ | IEEEGE[**2E**]FDE[**1E**]EEQ<br>**OR**<br>IEEEGE[**1E**]FDE[**2E**]EEQ |
| 4E | IEEEGE[**2E**]FDE[**2E**]EEQ | IEEEGE[**3E**]FDE[**1E**]EEQ<br>**OR**<br>IEEEGE[**2E**]FDE[**2E**]EEQ | IEEEGE[2**E**]FDE[**2E**]EEQ |
| 5E | IEEEGE[**3E**]FDE[**2E**]EEQ<br>**OR**<br>IEEEGE[**3E**]FDEE[**2E**]EQ | IEEEGE[**3E**]FDE[**2E**]EEQ | IEEEGE[3**E**]FDE[**2E**]EEQ |

**Table S4b.** Glutamylated peptides identified for tyrosinated β-tubulin (second biological replicate)

| **Number of E** | **Replicate 1** | **Replicate 2** | **Replicate 3** |
|---|---|---|---|
| 1E | IEEEGE[**1E**]FDEEEQY | IEEEGE[**1E**]FDEEEQY | IEEEGE[**1E**]FDEEEQY |
| 2E | IEEEGE[**1E**]FDE[**1E**]EEQY | IEEEGE[**1E**]FDE[**1E**]EEQY | IEEEGE[**1E**]FDE[**1E**]EEQY |
| 3E | IEEEGE[**2E**]FDE[**1E**]EEQY | IEEEGE[**2E**]FDE[**1E**]EEQY | IEEEGE[**2E**]FDE[**1E**]EEQY<br>**OR**<br>IE[**2E**]EEGEFDE[**1E**]EEQY |
| 4E | IEEEGE[**2E**]FDE[**2E**]EEQY | IEEEGE[**2E**]FDE[**2E**]EEQY | IEEEGE[**2E**]FDE[**2E**]EEQY |

**Table S5.** List of synthetic peptides and correction factors applied to corresponding peptides (500 fmol injection)

| Peptide | No of E | Correction factor |
|---|---|---|
| LEKDYEEVGAESADMDGEEDVEE* | 0 | 0.99±0.05 |
| LEKDYEEVGAESADMDGEEDVEE**Y*** | | 2.76±0.32 |
| LEKDYEEVGAESADMDGEEEDVEE* | 1 | 1 |
| LEKDYEEVGAESADMDGEEEDVEE**Y*** | | 2.16±0.12 |
| LEKDYEEVGAESADMDGEEEEDVEE | 2 | 1.59 |
| LEKDYEEVGAESADMDGEEEEDVEE**Y** | | 2.92 |
| LEKDYEEVGAESADMDGEEEEEDVEE* | 3 | 3.84±0.11 |
| LEKDYEEVGAESADMDGEEEEEDVEE**Y*** | | 4.56±0.56 |
| LEKDYEEVGAESADMDGEEEEEEDVEE | 4 | 5.28 |
| LEKDYEEVGAESADMDGEEEEEEDVEE**Y** | | 5.84 |
| LEKDYEEVGAESADMDGEEEEEEEDVEE* | 5 | 8.44±0.46 |
| LEKDYEEVGAESADMDGEEEEEEEDVEE**Y*** | | 8.29±0.37 |
| LEKDYEEVGAESADMDGEEEEEEEEDVEE | 6 | 9.76 |
| LEKDYEEVGAESADMDGEEEEEEEEDVEE**Y** | | 9.81 |
| LEKDYEEVGAESADMDGEEEEEEEEEDVEE | 7 | 11.57 |
| LEKDYEEVGAESADMDGEEEEEEEEEDVEE**Y** | | 12.01 |
| LEKDYEEVGAESADMDGEEEEEEEEEEDVEE | 8 | 14.21 |
| LEKDYEEVGAESADMDGEEEEEEEEEEDVEE**Y** | | 15.47 |
| LEKDYEEVGAESADMDGEEEEEEEEEEEDVEE | 9 | 18.39 |
| LEKDYEEVGAESADMDGEEEEEEEEEEEDVEE**Y** | | 21.74 |
| LEKDYEEVGAESADMDGEEEEEEEEEEEEDVEE* | 10 | 26.06±1.5 |
| LEKDYEEVGAESADMDGEEEEEEEEEEEEDVEE**Y*** | | 36.61±2.72 |
| IEEEGEFDEEEQ* | 0 | 1.47±0.1 |
| IEEEGEFDEEEQ**Y*** | | 5.85±0.49 |
| IEEEGEFDEEEEQ* | 1 | 0.93±0.08 |
| IEEEGEFDEEEEQ**Y*** | | 4.06±0.12 |
| IEEEGEFDEEEEEQ | 2 | 0.98 |
| IEEEGEFDEEEEEQ**Y** | | 2.87 |
| IEEEGEFDEEEEEEQ* | 3 | 1.03±0.03 |
| IEEEGEFDEEEEEEQ**Y*** | | 2.22±0.12 |
| IEEEGEFDEEEEEEEQ | 4 | 1.45 |
| IEEEGEFDEEEEEEEQ**Y** | | 2.85 |
| IEEEGEFDEEEEEEEEQ* | 5 | 2.44±0.14 |
| IEEEGEFDEEEEEEEEQ**Y*** | | 3.87±0.17 |

*Measured synthetic peptides for which the correction factor was calculated based on triplicate measurement average ± s.d. For the rest, correction factor was calculated based on linear regression between neighbouring points as shown in Figure S1.

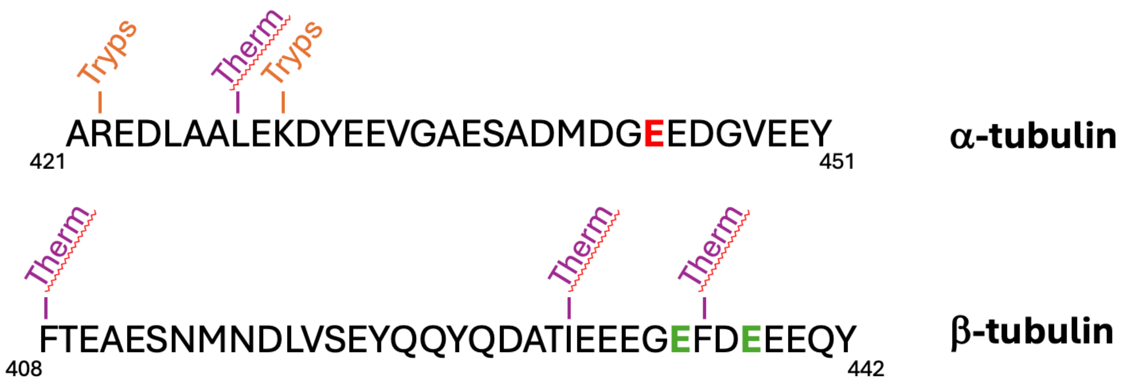


**Figure S1.** Predicted cleavage sites of α- and β-tubulin C-terminus for trypsin (Tryps), AspN and thermolysin (Therm).

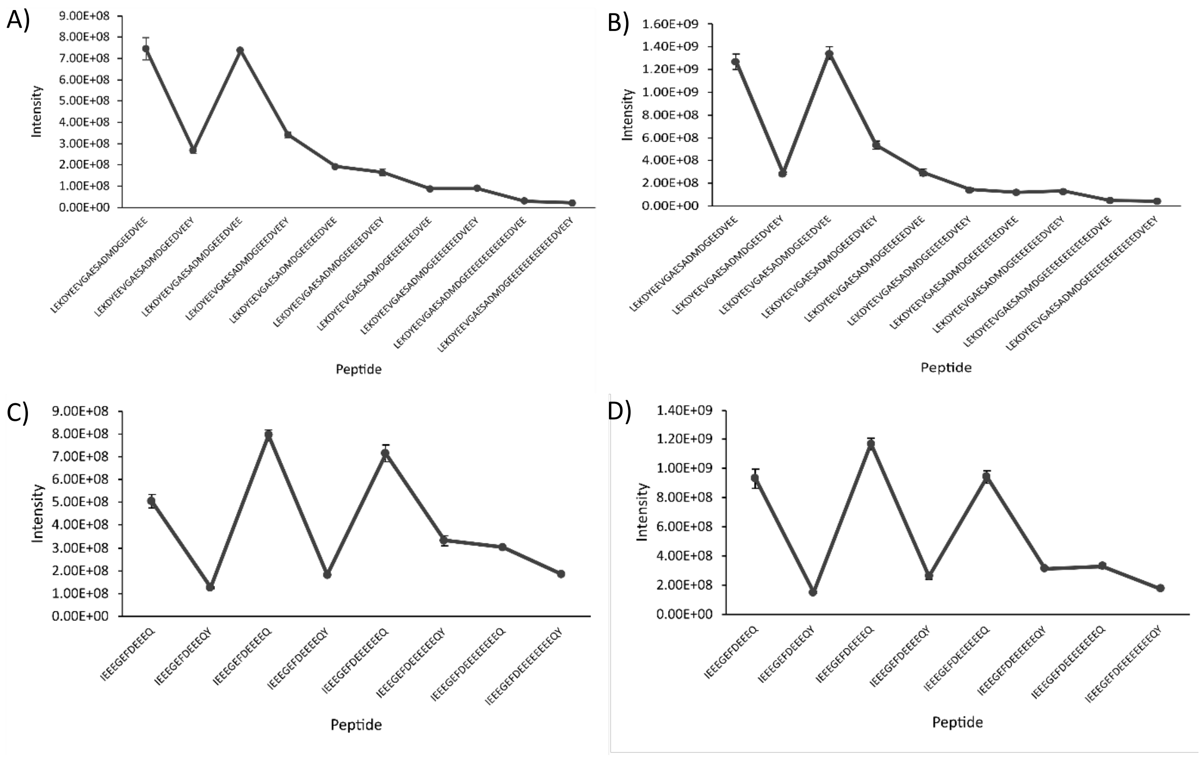


**Figure S2.** Synthetic peptide MS intensity response corresponding to alpha- (A-B) and beta- (C-D) tubulin C-terminals recorded at 500 fmol injection (left) or 225 fmol injection spiked in yeast digest (right).

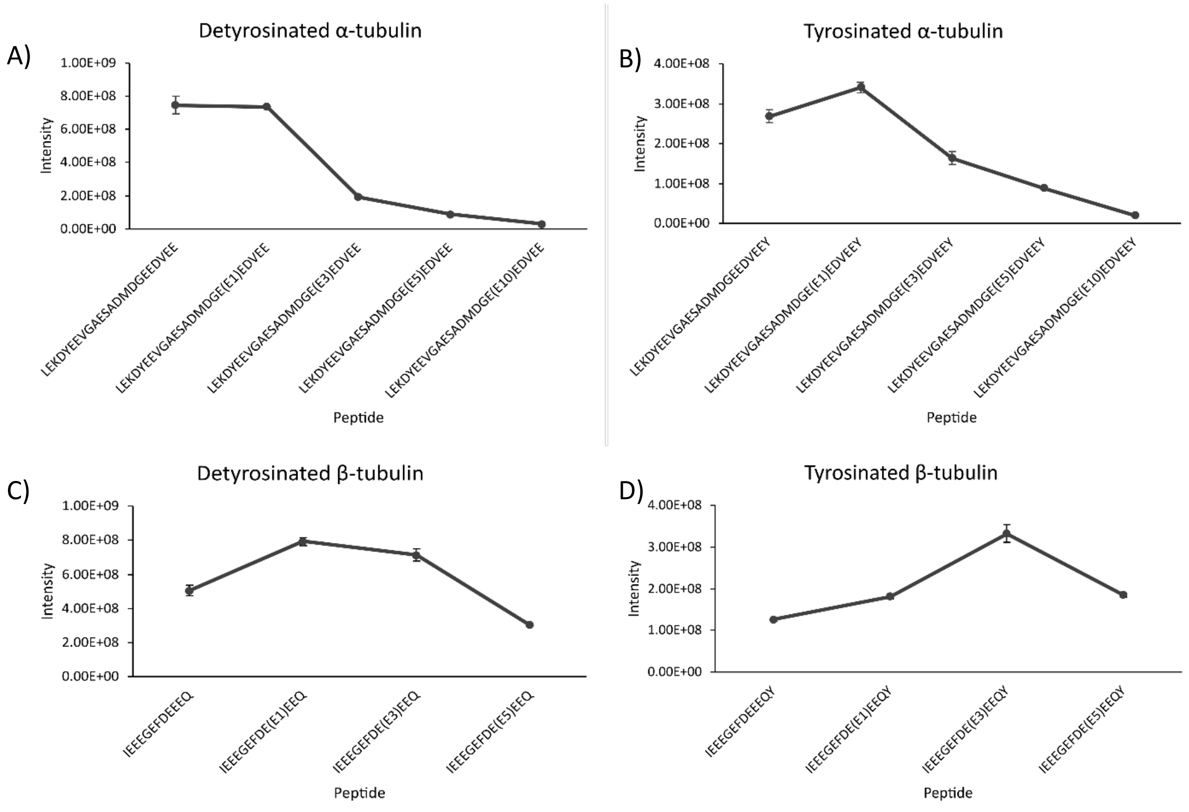


**Figure S3.** Synthetic peptide (500 fmol injection) MS intensity response corresponding to alpha- (A-B) or beta- (C-D) tubulin C-terminals in their detyrosinated (left) and tyrosinated (right) forms. Values for missing peptides with 2, 4, 6-10 for α- and 2, 4 E for β-tubulin, were approximated based on linear regression between neighbouring points for each.

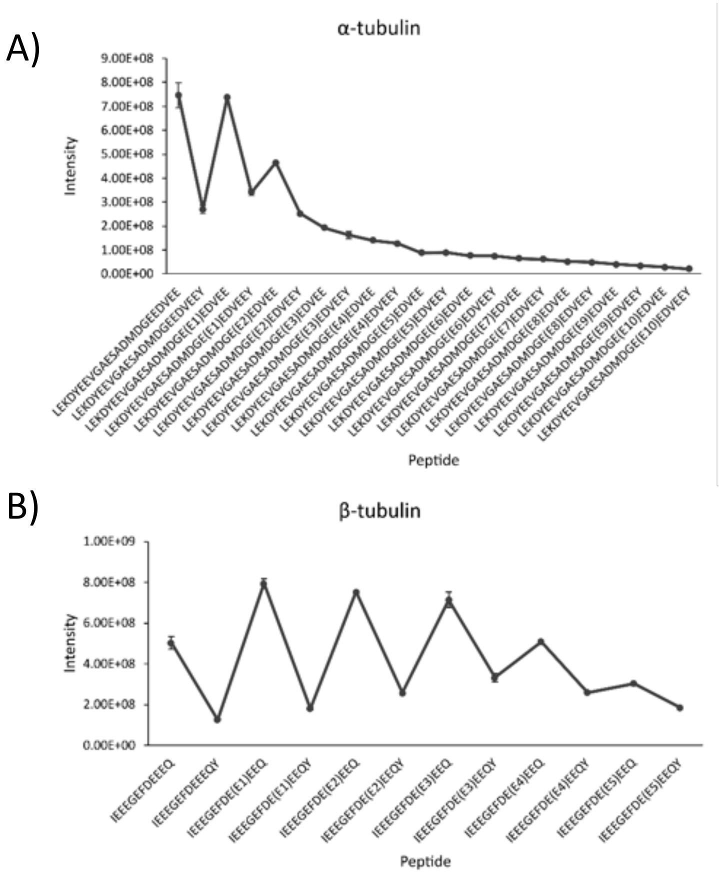


**Figure S4.** Synthetic peptide (500 fmol injection) MS intensity response corresponding to A) alpha- or B) beta-tubulin C-terminal ones, together with approximated values for missing peptides with 2, 4, 6-10 for α- and 2, 4 E for β-tubulin.

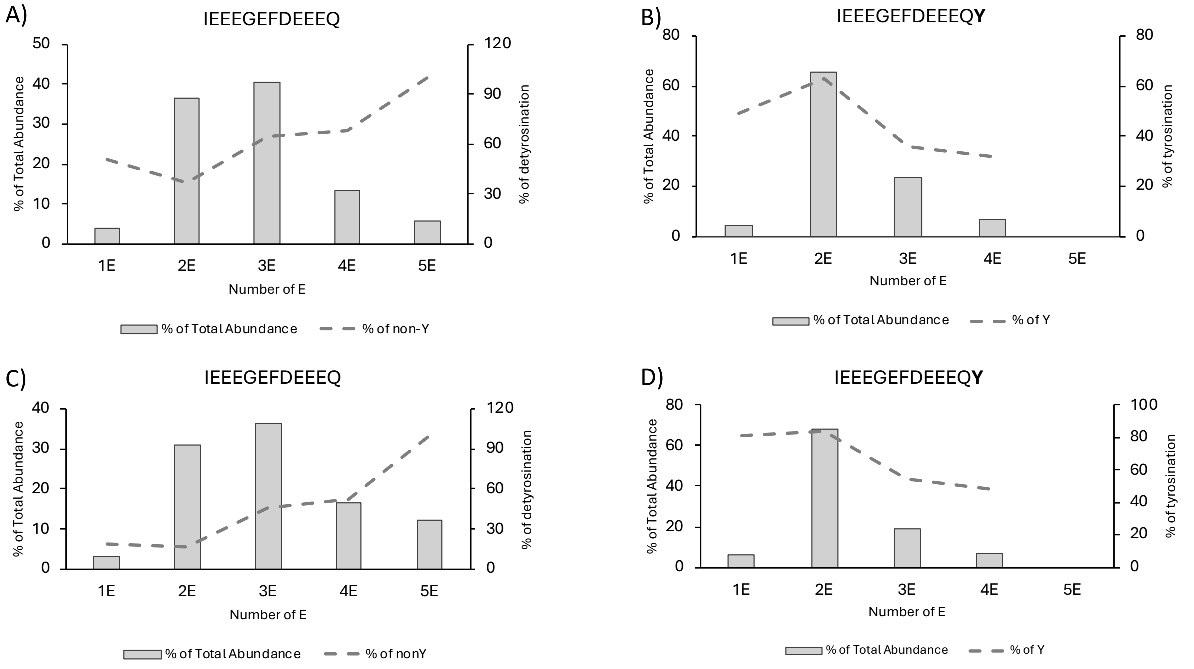

**Figure S5.** Quantification of β-tubulin C-terminal peptide variants identified in the first biological replicate: A) detyrosinated before correction B) tyrosinated before correction C) detyrosinated after correction D) tyrosinated after correction. Dashed lines represent the variation in detyrosination (A and C) or tyrosination (B and D) with increasing the number of supplementary glutamates.

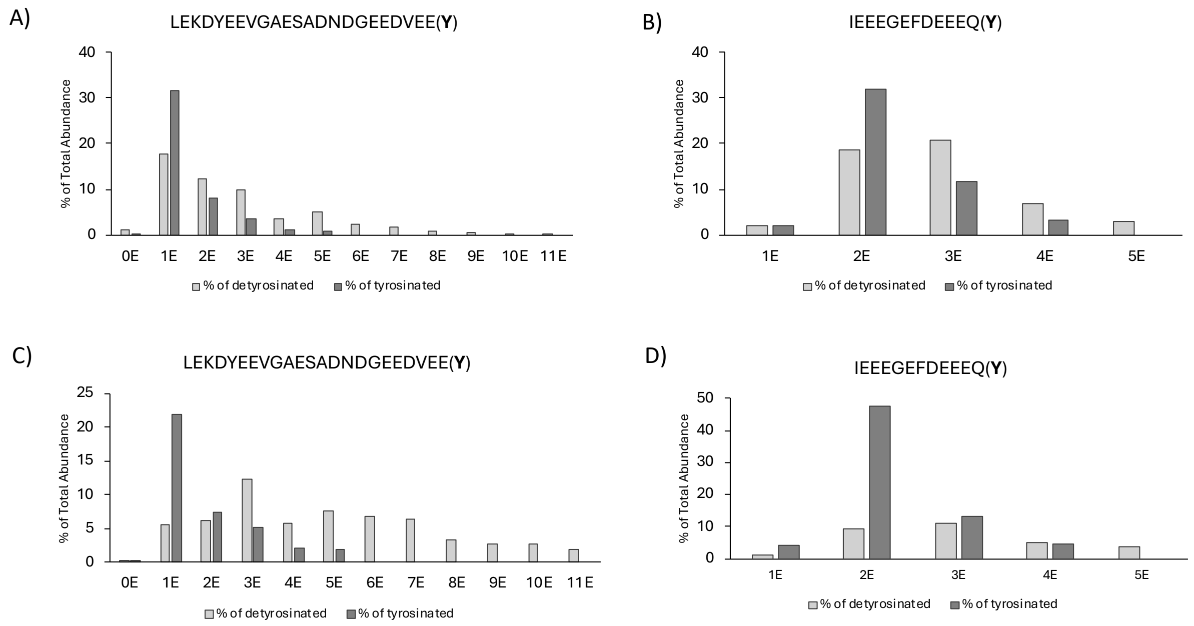


**Figure S6.** Relative abundances of detyrosinated and tyrosinated (poly)glutamylated peptide variants of α-tubulin A) before and C) after correction and β-tubulin B) before and D) after correction with respect to their total abundance.

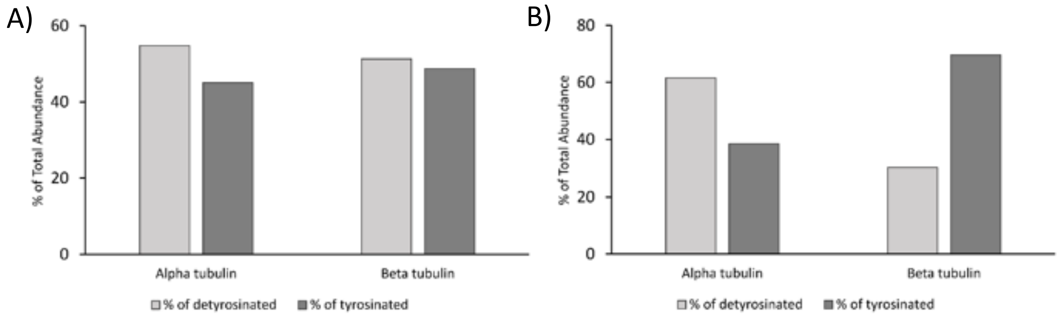


**Figure S7.** Relative abundances of detyrosinated and tyrosinated α- and β-tubulin in the first biological replicate A) before and B) after correction.

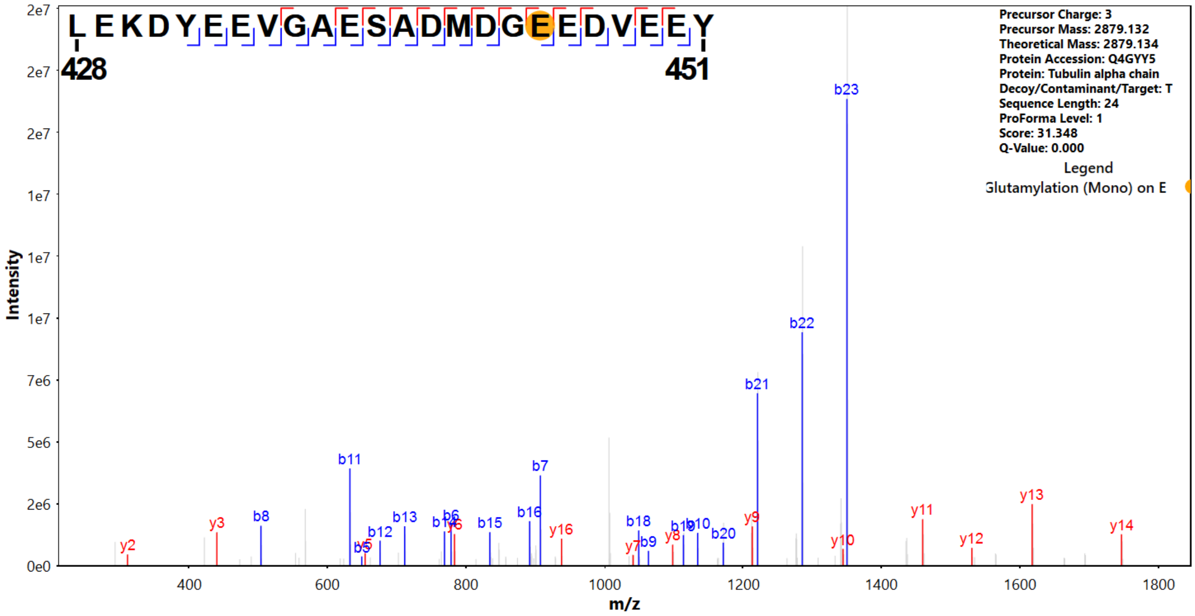


**Figure S8.** MS/MS spectrum of wild type flagella α-tubulin $_{428}$LEKDYEEVGAESADMDG**E**(E1)EDVEE**Y**$_{451}$ peptide.

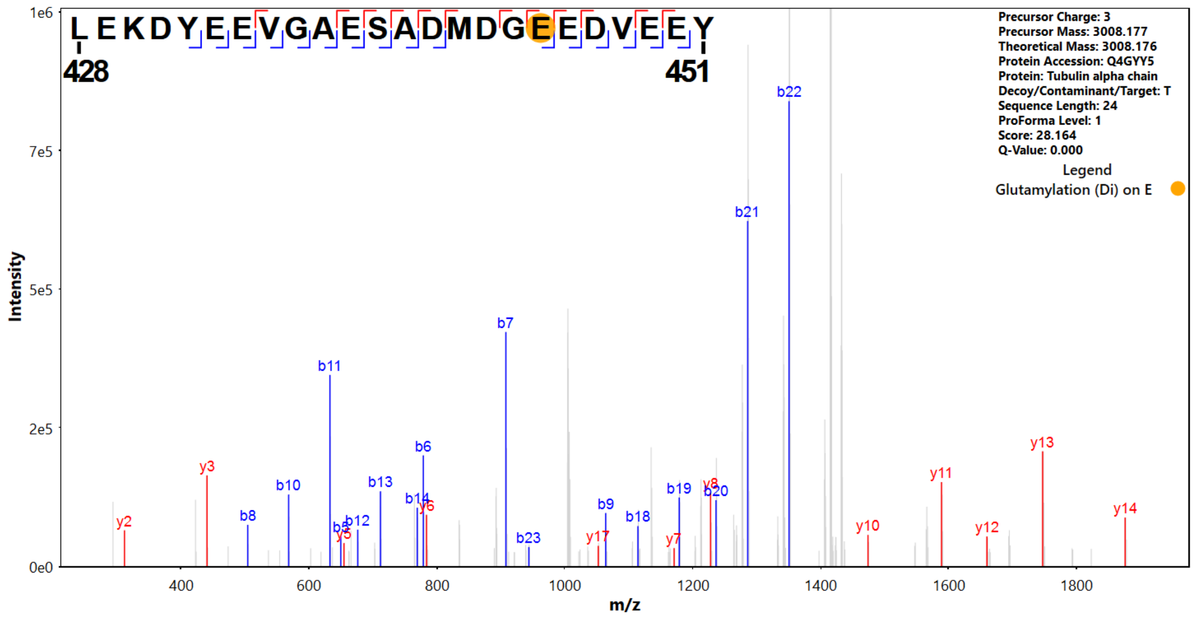


**Figure S9.** MS/MS spectrum of wild type flagella α-tubulin $_{428}$LEKDYEEVGAESADMDG**E**(E2)EDVEE**Y**$_{451}$ peptide.

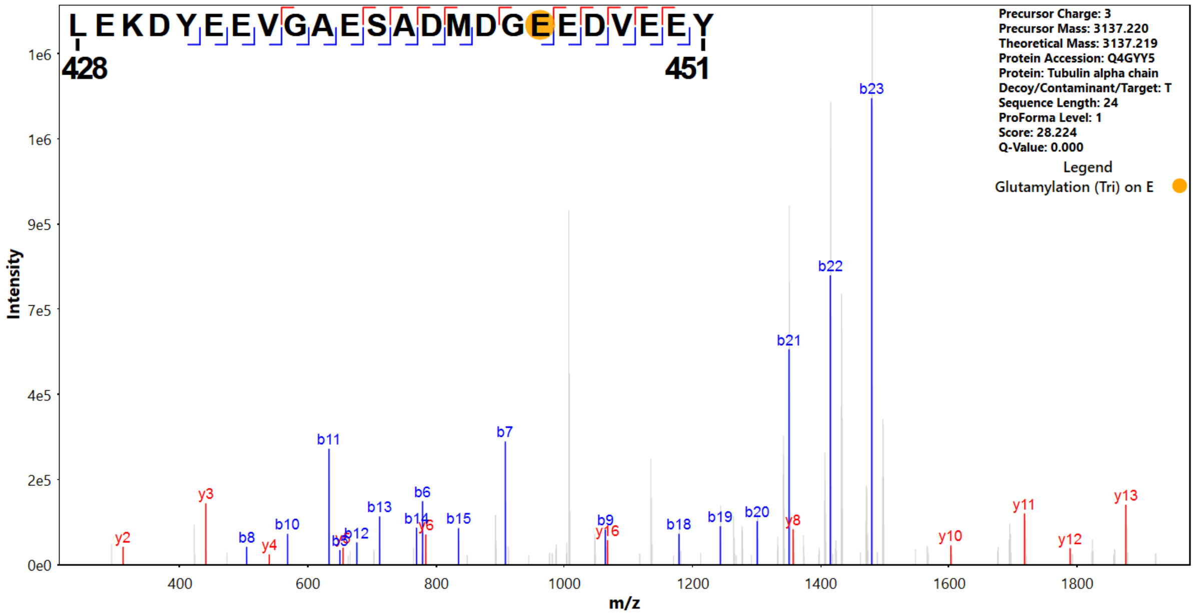


**Figure S10.** MS/MS spectrum of wild type flagella α-tubulin $_{428}$LEKDYEEVGAESADMDG**E**(E3)EDVEE**Y**$_{451}$ peptide.

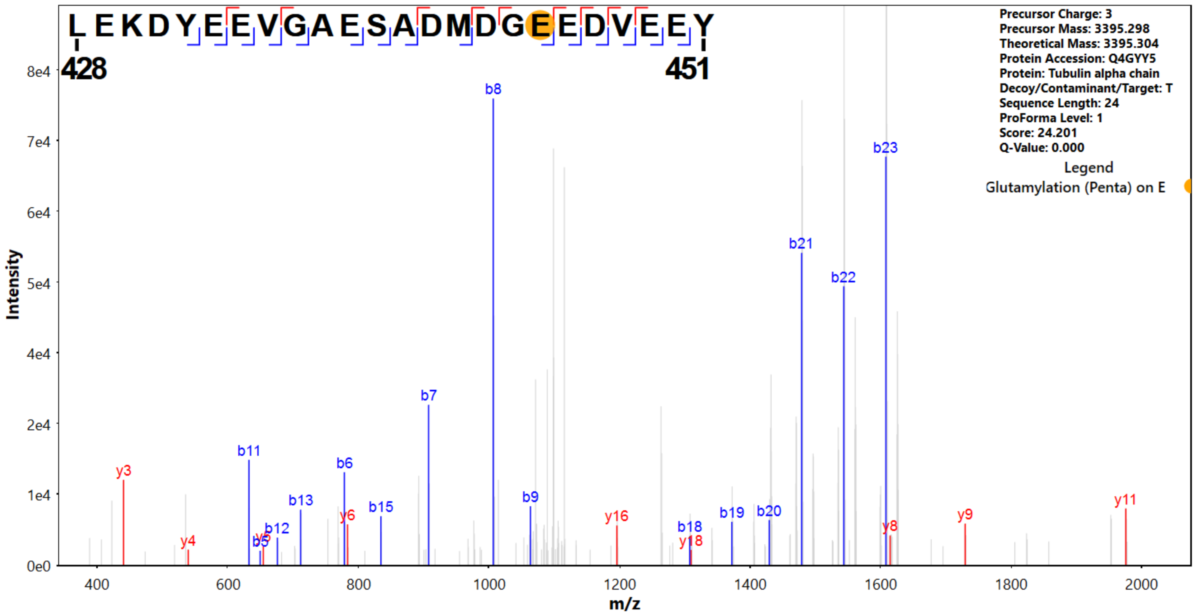


**Figure S11.** MS/MS spectrum of wild type flagella α-tubulin $_{428}$LEKDYEEVGAESADMDG**<u>E</u>**(E5)EDVEE**Y**$_{451}$ peptide.

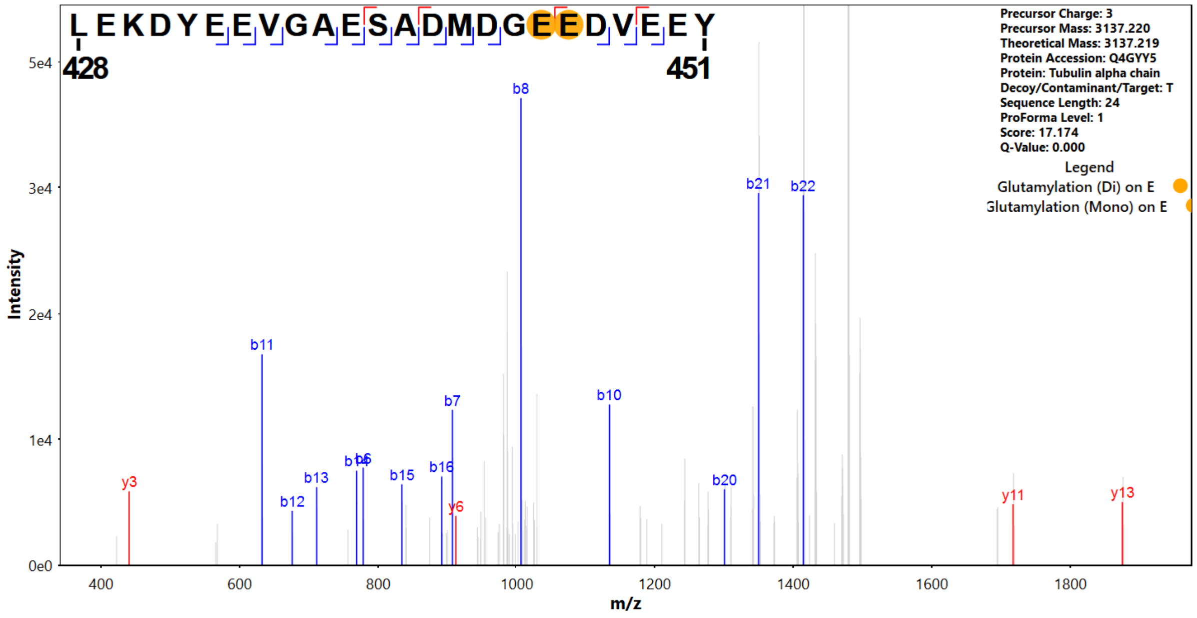


**Figure S12.** MS/MS spectrum of wild type flagella α-tubulin $_{428}$LEKDYEEVGAESADMDGE(E2)E(E1)DVEE**Y**$_{451}$ peptide.

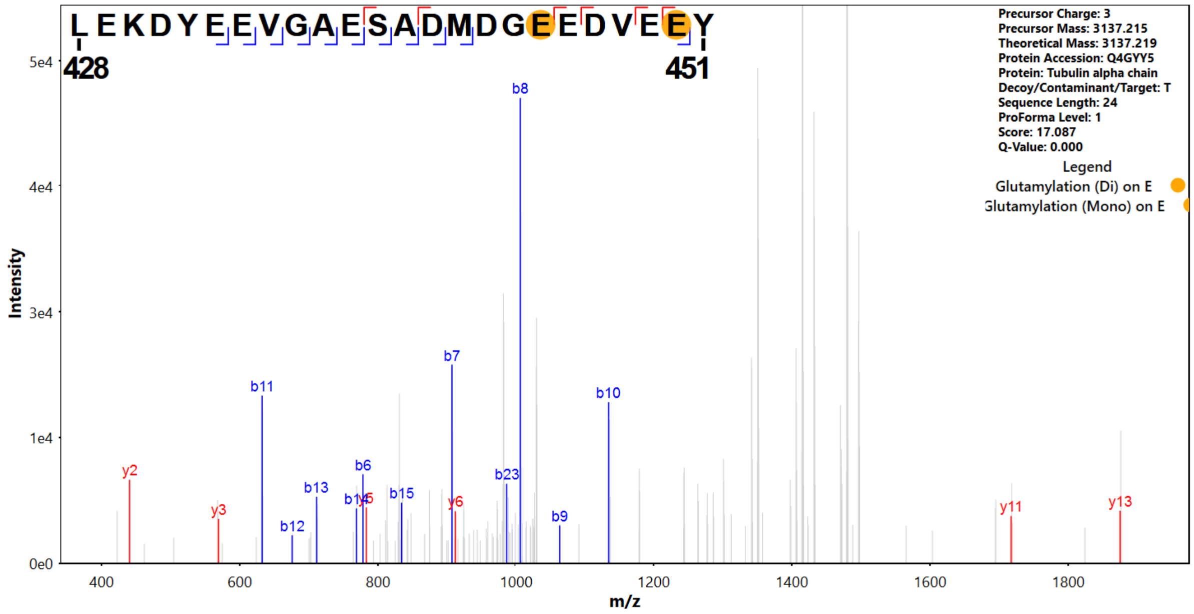


**Figure S13.** MS/MS spectrum of wild type flagella α-tubulin $_{428}$LEKDYEEVGAESADMDG**E**(E2)EDVE**E**(E1)**Y**$_{451}$ peptide.

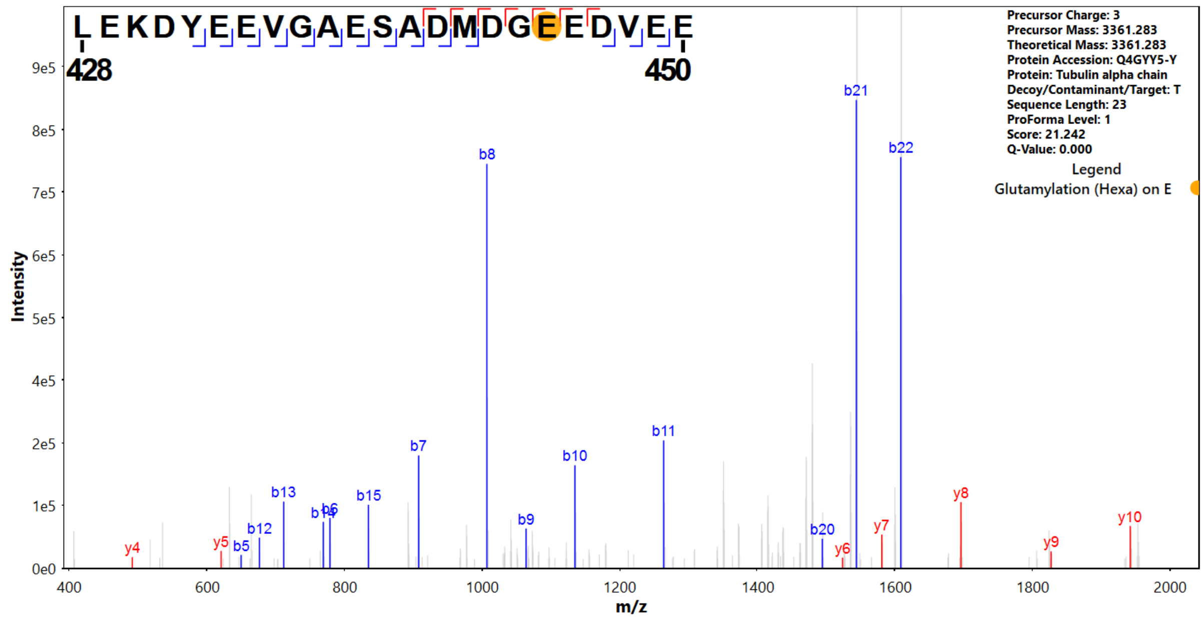


**Figure S14.** MS/MS spectrum of wild type flagella α-tubulin $_{428}$LEKDYEEVGAESADMDG**E**(E6)EDVEE$_{450}$ peptide.

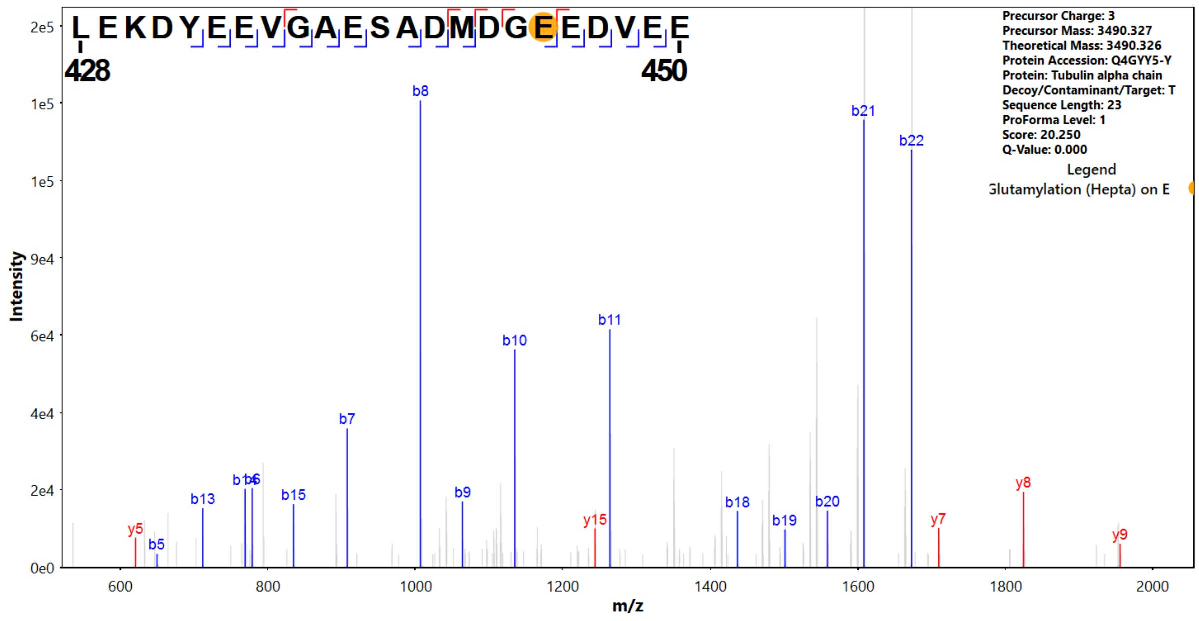


**Figure S15.** MS/MS spectrum of wild type flagella α-tubulin $_{428}$LEKDYEEVGAESADMDG**E**(E7)EDVEE$_{450}$ peptide.

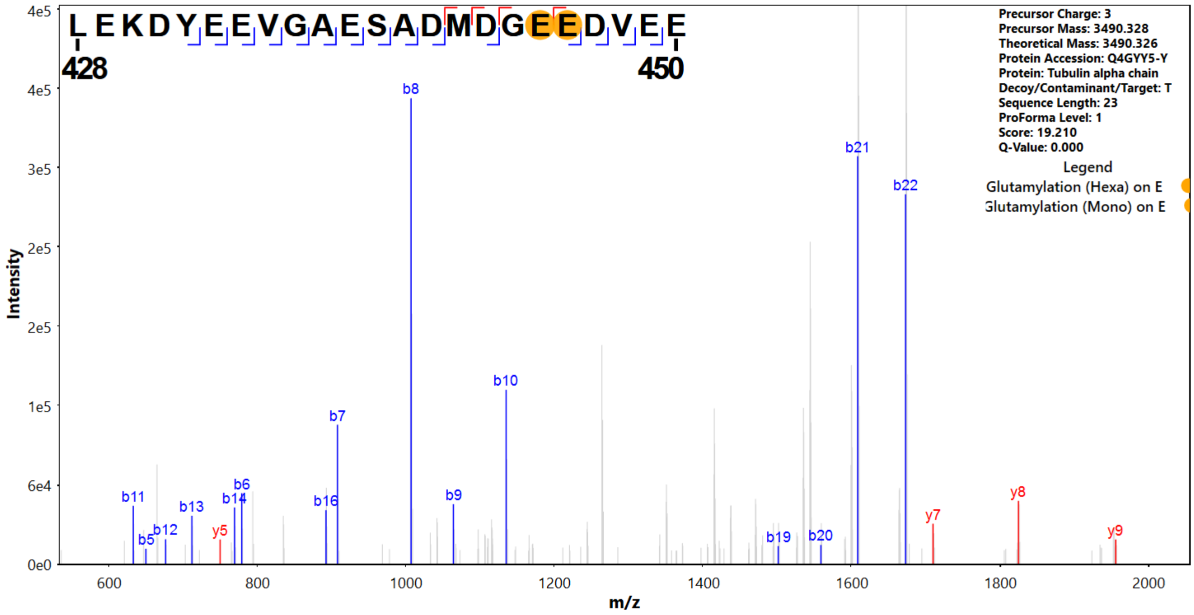


**Figure S16.** MS/MS spectrum of wild type flagella α-tubulin $_{428}$LEKDYEEVGAESADMDG**E**(E6)**E**(E1)DVEE$_{450}$ peptide.

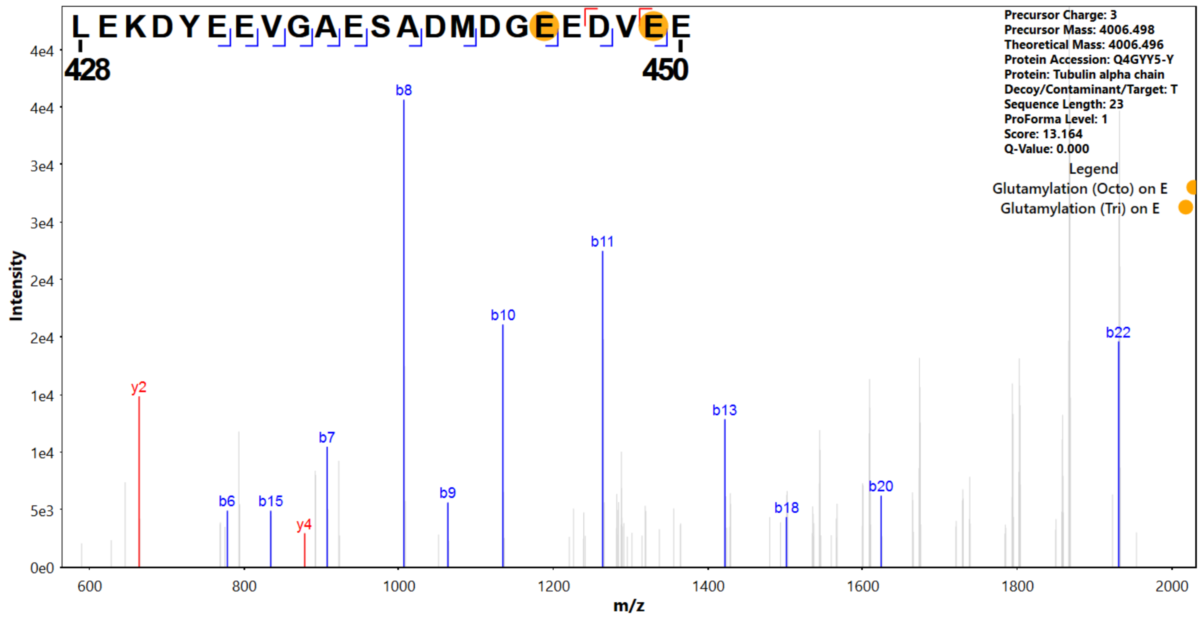


**Figure S17.** MS/MS spectrum of wild type flagella α-tubulin $_{428}$LEKDYEEVGAESADMDG**E**(E8)EDV**E**(E3)E$_{450}$ peptide.

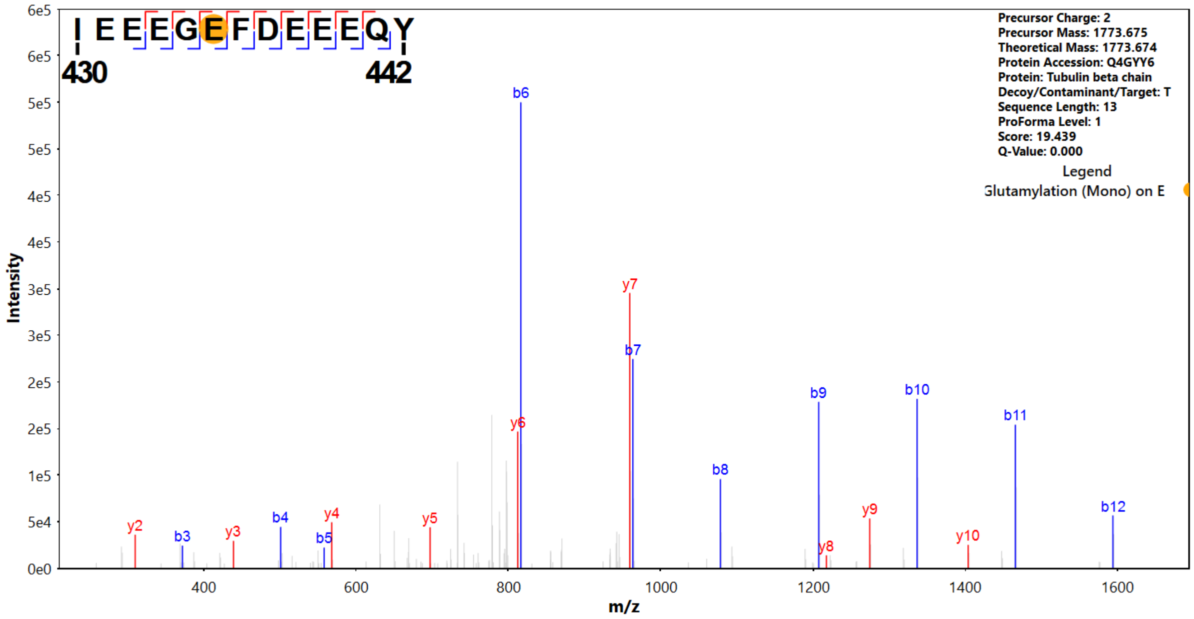


**Figure S18.** MS/MS spectrum of wild type flagella β-tubulin $_{430}$IEEEG**E**(E1)FDEEEQ**Y**$_{442}$ peptide.

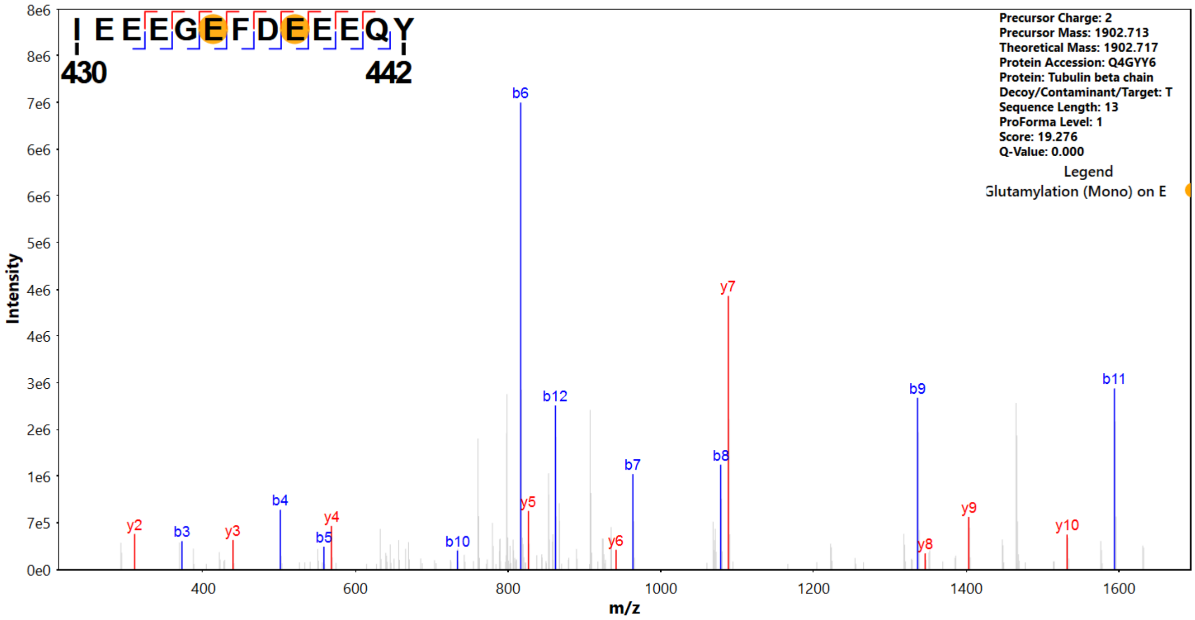


**Figure S19.** MS/MS spectrum of wild type flagella β-tubulin $_{430}$IEEEG**E**(E1)FD**E**(E1)EEQ**Y**$_{442}$ peptide.

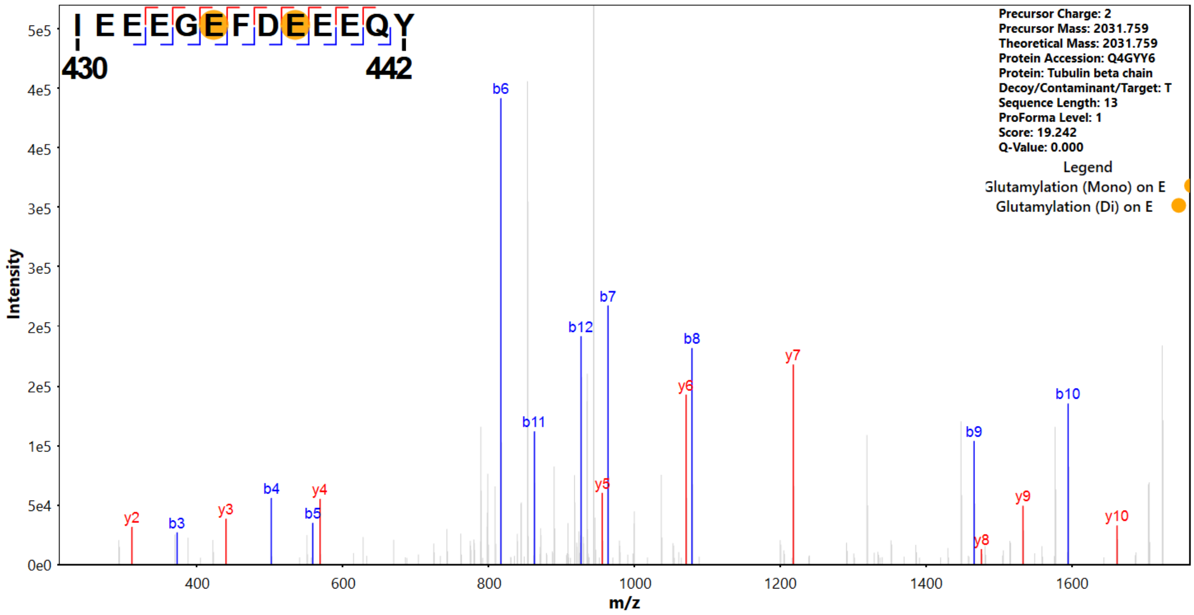


**Figure S20.** MS/MS spectrum of wild type flagella β-tubulin $_{430}$IEEEG**E**(E1)FD**E**(E2)EEQ**Y**$_{442}$ peptide.

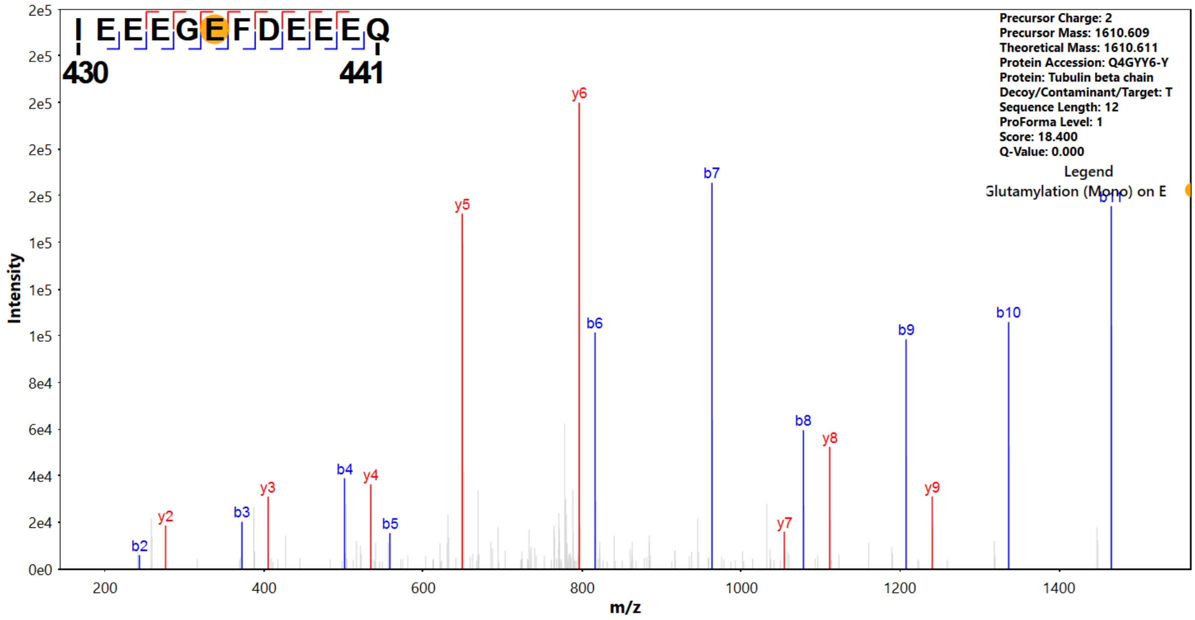


**Figure S21.** MS/MS spectrum of wild type flagella β-tubulin $_{430}$IEEEG**E**(E1)FDEEEQ$_{441}$ peptide.

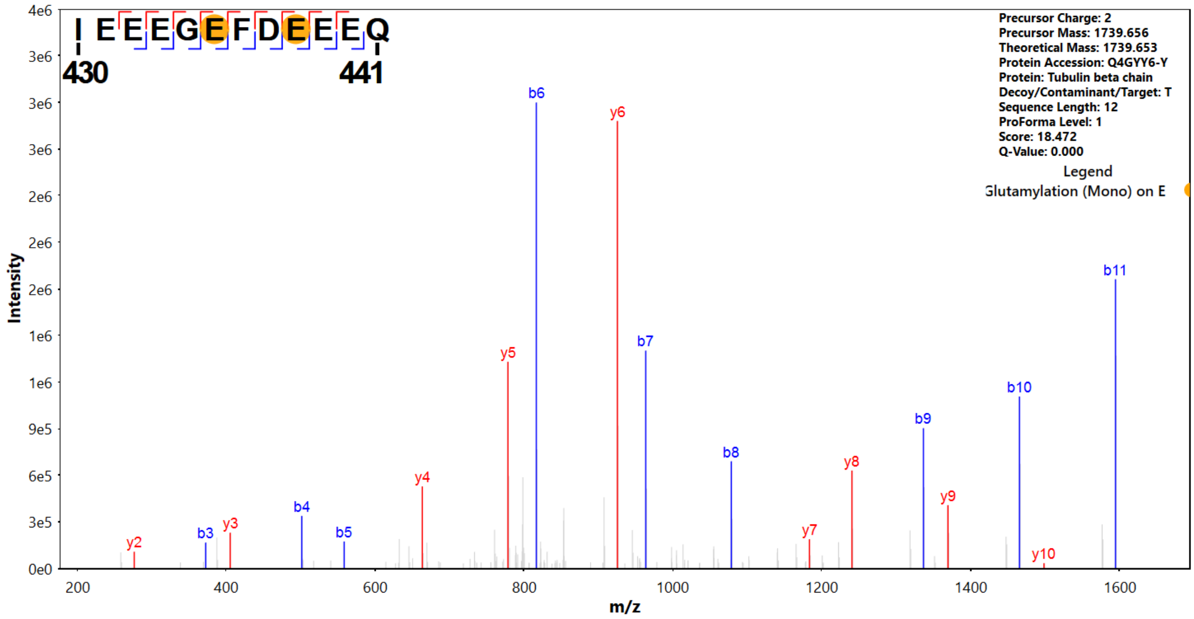


**Figure S22.** MS/MS spectrum of wild type flagella β-tubulin $_{430}$IEEEG**E**(E1)FD**E**(E1)EEQ$_{441}$ peptide.

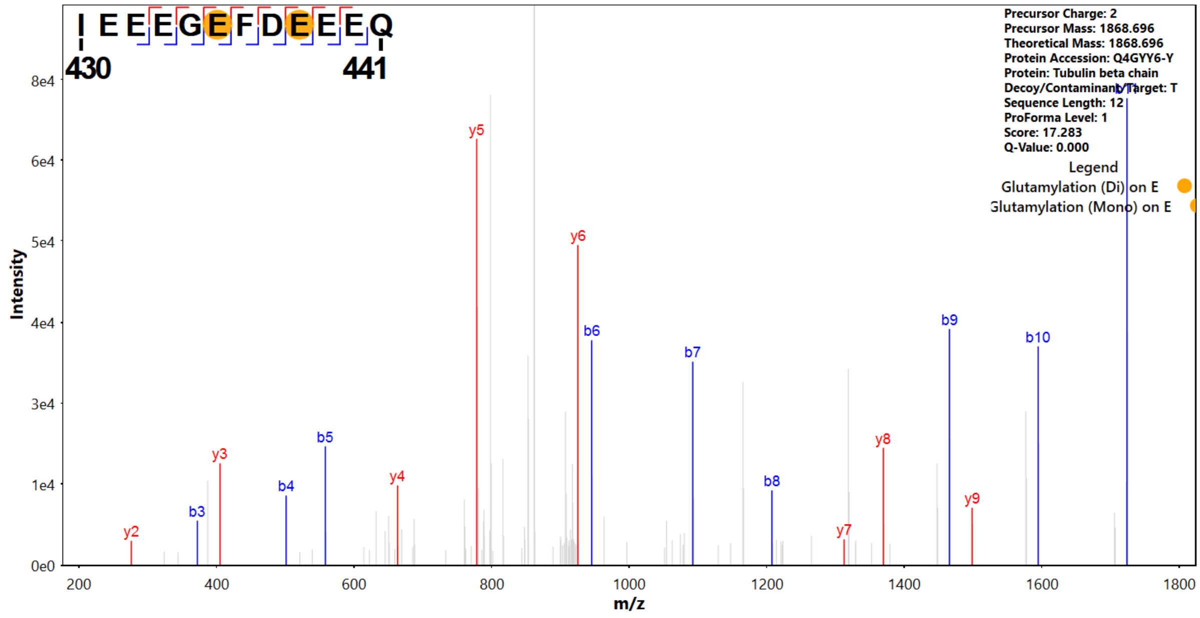


**Figure S23.** MS/MS spectrum of wild type flagella β-tubulin $_{430}$IEEEG<u>**E**</u>(E2)FD<u>**E**</u>(E1)EEQ$_{441}$ peptide.

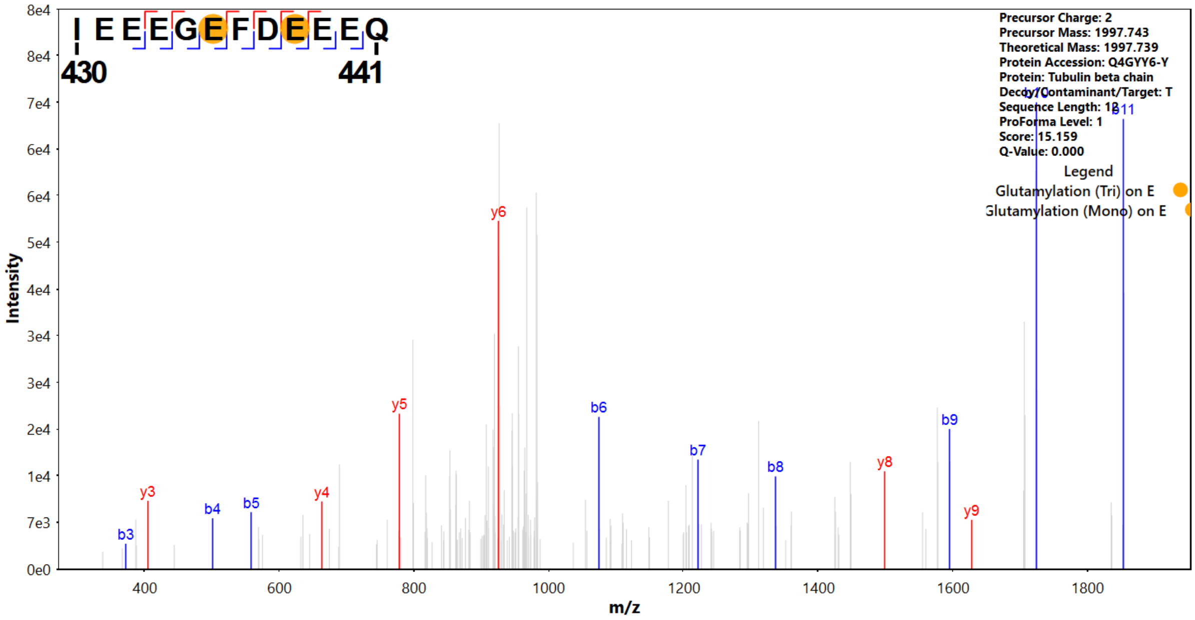


**Figure S24.** MS/MS spectrum of wild type flagella β-tubulin $_{430}$IEEEG**E**(E3)FD**E**(E1)EEQ$_{441}$ peptide.

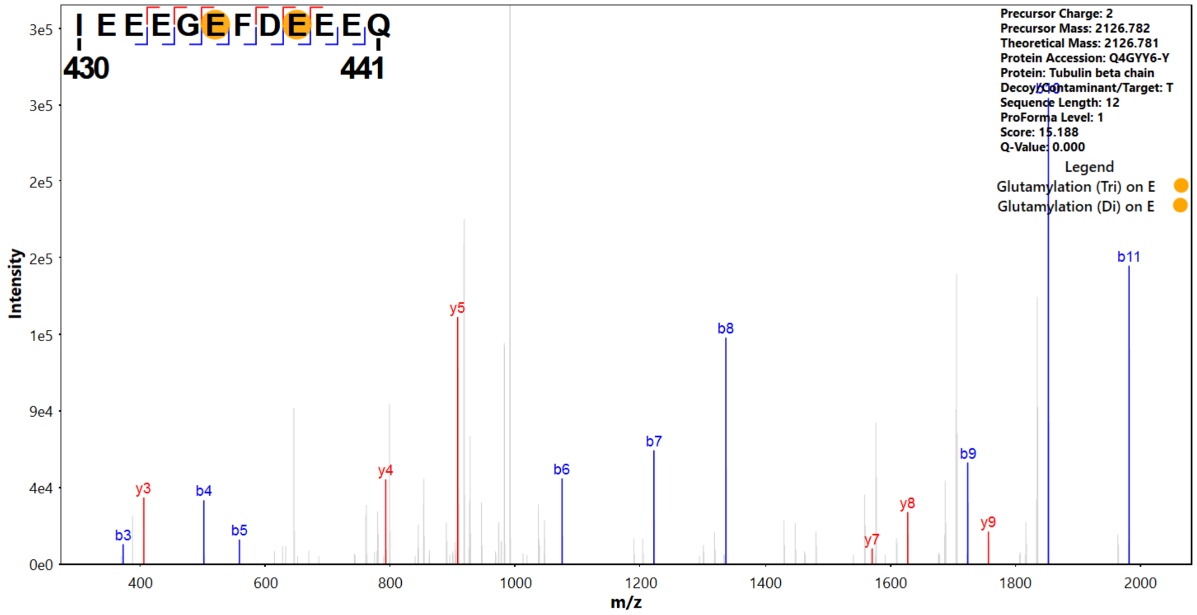


**Figure S25.** MS/MS spectrum of wild type flagella β-tubulin $_{430}$IEEEG**E**(E3)FD**E**(E2)EEQ$_{441}$ peptide.
